# Hippo-YAP1/TAZ-TEAD signaling coordinates myoblast fusion and sarcomere assembly in skeletal muscle development

**DOI:** 10.1101/2024.06.11.598443

**Authors:** Lea Gessler, Anna Siudzińska, Tomasz J. Prószyński, Marco Sandri, Björn von Eyss, Said Hashemolhosseini

**Affiliations:** Institute of Biochemistry, Medical Faculty, Friedrich-Alexander-University of Erlangen-Nürnberg, Erlangen, Germany; Łukasiewicz Research Network-PORT Polish Center for Technology Development, Wrocław, Poland; Department of Biomedical Science, Venetian Institute of Molecular Medicine, University of Padova, Padova, Italy; Transcriptional Control of Tissue Homeostasis Lab, Leibniz Institute on Aging, Fritz Lipmann Institute e.V., Jena, Germany

**Keywords:** Hippo signaling, YAP1 / TAZ, TEAD transcription factors, Skeletal muscle morphogenesis, Myoblast fusion, Sarcomere assembly

## Abstract

Hippo-YAP1/TAZ-TEAD signaling has been implicated in myogenesis, but how this pathway coordinates myoblast fusion with sarcomere assembly *in vivo* is poorly understood. Here we dissect the roles of YAP1 and TAZ in skeletal muscle development of mice using muscle-specific single and double knockouts combined with multi-omics and regulatory analysis. Muscle-specific double knockout of *Yap1* and *Wwtr1* (*Taz*) causes perinatal lethality with failure of myofiber maturation, characterized by secondary myotubes that remain arrested in small, doughnut-like configurations and by disorganized, incomplete sarcomeres. Developmental analysis reveals non-redundant functions of YAP1 and TAZ in coordinating primary and secondary myogenesis and establishing fiber-type architecture. Integrative RNA-seq, TEAD1/TEAD4 ChIP-seq and single-cell ATAC-seq identify a conserved YAP1/TAZ-TEAD transcriptional program that jointly regulates myoblast fusion genes (*Mymk*, *Mymx*) and sarcomere genes (*Myh3*, *Ttn*) via MCAT-containing regulatory elements. Together, these findings establish Hippo-YAP1/TAZ-TEAD signaling as a central coordinator of myoblast fusion and sarcomere assembly in skeletal muscle, providing a mechanistic framework for Hippo-dependent control of tissue morphogenesis.

**Key Points:**

• **YAP1 and TAZ have non-redundant roles in skeletal muscle development.**
Muscle-specific single knockouts reveal opposite effects on fiber size and fiber-type composition at adulthood, while double knockout causes perinatal lethality with profound failure of myofiber maturation.
• **Hippo-YAP1/TAZ signaling is essential for the transition from myoblast fusion to stable secondary myotubes.**
In double-knockout embryos, secondary myotubes form but remain arrested in small, doughnut-like configurations and regress in distal hindlimb muscles.
• **YAP1/TAZ-TEAD networks control sarcomere assembly and architecture *in vivo*.**
Neonatal double-knockout myofibers display disorganized and incomplete sarcomeres by histology and electron microscopy, indicating a direct requirement for Hippo signaling in contractile apparatus morphogenesis.
• **An integrated YAP1/TAZ-TEAD transcriptional program jointly regulates fusion genes and sarcomere genes.**
RNA-seq, TEAD1/TEAD4 binding, and chromatin accessibility analyses identify MCAT-containing regulatory elements in *Mymk*, *Mymx*, *Myh3*, *Myl1*, *Myl2* and *Ttn* that link Hippo signaling to both myoblast fusion and thick-filament gene expression.
• **MCAT-driven regulatory elements function as TEAD-responsive enhancers in myogenic cells.**
Reporter assays demonstrate increased activity of selected MCAT-containing elements in differentiating myotubes, supporting their role as functional nodes through which Hippo-YAP1/TAZ-TEAD networks orchestrate muscle fiber growth and sarcomere gene programs.

## INTRODUCTION

Skeletal muscle development in mammals proceeds through embryonic and fetal myogenesis, followed by postnatal growth and regeneration. In mice, a first wave of myofiber formation between embryonic days 10.5 and 12.5 (E10.5-E12.5) generates primary fibers from somite-derived myoblasts (1,2). These fibers enlarge rapidly, initially show a characteristic “doughnut”-shaped morphology in cross section, and provide an essential scaffold for later fiber formation. A second wave of myogenesis between E14.5 and E17.5 (secondary myogenesis) generates secondary myofibers as fetal myoblasts differentiate and fuse in close association with the pre-existing primary fibers. Embryonic myosin *Myh3* is expressed in early fibers of both waves, and the morphological distinction between primary and secondary fibers diminishes as development proceeds.

Postnatally, satellite cells become the principal source of myonuclei for muscle growth and regeneration. Upon activation, they proliferate and fuse with existing myofibers or with each other, thereby contributing new myonuclei to growing or regenerating fibers (3). In adult muscle, quiescent satellite cells reside between the basal lamina and the sarcolemma, express *Pax7*, and upregulate *Myod1* and *Myog* upon injury to drive efficient regeneration (3).

The Hippo pathway is a key regulator of organ size, tissue regeneration, and stem cell self-renewal (1). In mammals, activation of MST1/2 and LATS1/2 kinases leads to phosphorylation of the transcriptional coactivators YAP1 and TAZ (Wwtr1), thereby limiting their stability, nuclear localization, and transcriptional activity (2). In their active nuclear form, YAP1 and TAZ associate with TEA domain transcription factors TEAD1-4 to drive expression of target genes (4). TEAD1, originally identified as TEF-1, binds the M-CAT motif 5′-CATTCCT-3′, which is enriched in regulatory regions of many muscle-specific genes (5–7). Genome-wide studies have shown that YAP1 and TAZ bind TEADs at promoters and distal enhancers, where they cooperate with other transcriptional regulators such as AP-1 (8–11). TEAD activity is further modulated by VGLL proteins, with VGLL1-3 acting as tissue-restricted coactivators and VGLL4 competing with YAP1 as an inhibitor of TEAD-dependent transcription (12–16). Although YAP1 and TAZ often display overlapping activity toward TEAD transcription factors (17), genetic studies have revealed non-redundant functions. Germline *Yap1* knockout is embryonic lethal, whereas *Wwtr1* knockout mice can survive to adulthood but display defects in selected organs (18–20). Comparative analyses of *Yap1*, *Wwtr1*, and *Yap1*/*Wwtr1* double knockout cells showed that loss of *Yap1* has a stronger impact on cell size, metabolism, proliferation, and transcriptomic deregulation than loss of *Wwtr1*, and that the double knockout more closely resembles Yap1 single knockout (21). In skeletal muscle, several studies have suggested divergent roles of YAP1 and TAZ in the myogenic lineage and in the control of fiber size and metabolism (22–25).

Hippo and canonical Wnt/β-catenin (CTNNB1) pathways display multiple points of crosstalk (26). Cytoplasmic YAP1 and TAZ can restrain canonical Wnt signaling by interfering with DVL phosphorylation, sequestering CTNNB1, and participating in the CTNNB1 destruction complex (27–30). Conversely, in Wnt-ON states, dissociation of YAP1/TAZ from the destruction complex allows CTNNB1 and YAP1/TAZ to accumulate in the nucleus and activate their respective transcriptional programs (27,29–32). In skeletal muscle, we previously showed that canonical Wnt signals enhance TEAD-dependent reporter activity during myoblast differentiation, indicating that YAP1/TAZ-TEAD signaling is engaged in this context (33).

TEAD factors themselves are required for proper muscle differentiation. Combined knockdown of Tead1-4 in primary myoblasts or C2C12 cells severely impairs differentiation, and selective loss of Tead1 or Tead4 reduces myotube formation *in vitro* (34). Chromatin profiling revealed dynamic TEAD1 and TEAD4 occupancy at genes controlling Hippo, Wnt, TGFβ, and myogenic programs (34), and single-nucleus multi-omic analyses identified a transcriptional module involving MYOG, KLF5, and TEAD4 that drives the transition from developmental to mature myofibers (35). These observations implicate TEAD factors and their coactivators as central regulators of myogenic gene expression.

We and others have begun to dissect YAP1/TAZ functions in skeletal muscle fibers *in vivo*. Muscle-specific deletion of *Yap1* or *Wwtr1* individually results in reduced grip strength and fragmented neuromuscular junctions (NMJs), whereas muscle-specific *Yap1*/*Wwtr1* conditional double knockout (cDKO) mice die at birth, almost completely lack NMJs, and exhibit severely abnormal endplate morphology and innervation (23,36). Preliminary findings on structural muscle defects in Yap1/Wwtr1 double-knockout mice were previously made available as a preprint (37); the present study substantially extends this work through embryonic developmental staging, cell-intrinsic fusion assays, and functional enhancer validation, as detailed below. *In silico* analysis of TEAD1 and TEAD4 binding sites identified evolutionarily conserved TEAD motifs in key synaptic genes that lie within open chromatin and drive reporter activity in muscle cells (23). Loss of *Yap1* also impairs reinnervation after denervation, at least in part via reduced CTNNB1 activity (36). Despite these insights into synaptic function, the roles of YAP1 and TAZ in embryonic and fetal myofiber formation and in sarcomere gene regulation *in vivo* remain poorly understood.

The present study aimed to define the roles of YAP1 and TAZ in the development, structure, and function of skeletal muscle fibers and to understand how Hippo-YAP1/TAZ-TEAD networks coordinately regulates myoblast fusion and sarcomere gene programs. Using muscle-specific deletions of *Yap1*, *Wwtr1*, or both, we show that single-muscle knockouts are viable but exhibit altered expression of myogenic genes accompanied by changes in fiber-type distribution and fiber size. In contrast, cDKO mice die perinatally and display defective myotube formation with regression of distal hindlimb muscles and pronounced sarcomere abnormalities in neonatal myofibers. By integrating RNA-seq, ChIP-seq, scATAC-seq, evolutionary conservation analysis, and reporter assays, we identify a YAP1/TAZ-TEAD transcriptional program that directly engages key fusion and structural genes, including Myomaker, myosin light and heavy chains, and Titin. These findings establish YAP1/TAZ-TEAD-mediated transcription as a fundamental regulator of myotube formation and sarcomere gene expression, providing a mechanistic framework for skeletal muscle morphogenesis and suggesting potential avenues for targeting Hippo signaling in muscle disease.

## MATERIALS AND METHODS

### Immunofluorescence staining and fluorescence microscopy

For histochemical and immunofluorescence analyses, gastrocnemius and soleus muscles from adult mutant and control mice were flash-frozen in 2-methylbutane cooled in liquid nitrogen. Embryos were dissected at the indicated developmental stages, and newborn pups were decapitated immediately after birth for immunofluorescence experiments. Samples were fixed overnight in 4% paraformaldehyde, cryoprotected in 15% and 30% sucrose, embedded in OCT and cryosectioned (10 µm for adult muscle, 14 µm for embryonic/neonatal tissue) from the mid-region of each muscle or limb; adjacent sections were used to detect MyH-emb, MyH-slow, and MyH-fast isoforms.

For immunofluorescence staining, tissue sections were fixed for 10 min in 2% PFA, permeabilized for 15 min in 0.1% Triton X-100 and 100 mM glycine, and blocked with M.O.M. Blocking Reagent (Vector Laboratories, Eching, Germany) for 1 h at room temperature as described previously (38). Sections were incubated overnight at 4 °C with primary antibodies (Supplementary Table 1), washed three times in PBS (10 min each), and incubated for 2 h at room temperature with appropriate secondary antibodies (39). After nuclear counterstaining with DAPI, sections were mounted in Mowiol mounting medium.

### Histochemical staining

Hematoxylin and eosin (H&E) staining: H&E staining was performed using Mayer’s hemalum and Eosin solutions (Merck) following standard procedures, and sections were dehydrated and mounted in DPX.

Cytochrome c oxidase (COX) staining: Sections were incubated for 60 min at 37 °C in a reaction buffer containing 50 mM phosphate buffer (pH 7.4), 3,3′-diaminobenzidine tetrahydrochloride (DAB; D8001, Sigma-Aldrich), catalase (20 mg/mL; S41168, Sigma-Aldrich), sucrose, and cytochrome c (C2037, Sigma-Aldrich). After incubation, samples were rinsed with water and mounted in DPX.

Alcian blue staining: Whole-mount skeletal staining of newborn pups was performed immediately after decapitation according to a published protocol (40).

### Myotube analysis

Primary myotubes were generated from control and *Yap1*/*Wwtr1^Δ^*^/Δ^::Pax7-CreER^T2^ skeletal muscle progenitor cells after 4-hydroxytamoxifen (4-OHT)-induced recombination in culture. Satellite cells were isolated as previously described (23) and differentiated for 6 days. Intact myotubes were visualized by fixation in 2% PFA and incubation with Rhodamine Fluor-conjugated bungarotoxin (BTX; 1:2500).

### Microscopy and image analysis

Fluorescence and bright-field images were acquired using a Zeiss Axio Examiner Z1 microscope with AxioCam cameras and ZEN Blue software and analyzed with ZEN Blue and ImageJ. Whole-mount images of hindlimb regions and of the hindlimb body part of neonatal mutant and control mice were obtained using a Zeiss Discovery V8 stereomicroscope equipped with an AxioCam HRm camera. Image analysis of whole-mount preparations was performed using ZEN Blue software.

### Electron microscopy

For ultrastructural analysis, neonatal diaphragms were dissected, fixed in 4% PFA, and stained en bloc using a modified OTO protocol (41). Samples were rinsed with PBS, stained with 2% OsO₄, treated with 2.5% potassium ferrocyanide, washed, and incubated with thiocarbohydrazide at 40 °C. After restaining with 2% OsO₄ and washing, samples were incubated overnight at 4 °C in 1% uranyl acetate. The following day, samples were warmed to 50 °C, washed with ultrapure water, dehydrated through a graded ethanol series, and embedded in Agar Low Viscosity Resin (AGR1078, Agar Scientific). Ultrathin sections were cut with a diamond knife, mounted on copper grids, and imaged using a scanning transmission electron microscopy (STEM) detector in a Zeiss Auriga SEM/FIB microscope operating at an accelerating voltage of 20 kV.

### In silico analysis of putative Tead binding sites

For *in silico* analysis of TEAD binding sites in myogenic genes, TEAD1 and TEAD4 ChIP-seq data from C2C12 cells at day 0 and day 6 of differentiation (34) were screened for occupancy in genomic regions encompassing sarcomere genes (Myh3, Myl1, Myl2, Ttn) and myogenic genes (Mymk, Mymx, Myod1, Myog), bounded by their respective 5′ and 3′ neighbouring genes. The UCSC Genome Browser (mouse July 2007 assembly, NCBI37/mm9) was used to identify these regions and export genomic sequences in FASTA format. Sequences were scanned in the JASPAR 2018 database (http://jaspar.genereg.net) (42) using the TEAD4 matrix model (MA0809.1) with the recommended relative profile score threshold of 80%. The resulting predicted TEAD4 binding sites were mapped back to chromosomal coordinates, and sites listed in Supplementary Table 3 showed high basewise conservation among mammalian species, as indicated by positive PhyloP scores (43). scATAC-seq datasets were examined for open chromatin regions in the same genes using CATLAS (https://decoder-genetics.wustl.edu/catlasv1/catlas_humanenhancer/#!/).

### Plasmids and Luciferase assay

Genomic regions of interest were amplified using primers containing KpnI or XhoI restriction sites (see Suppl. Table S4) to enable directional cloning into the pGL2-basic luciferase reporter vector (a gift from Michael Wegner, Friedrich-Alexander-Universität Erlangen-Nürnberg, Germany). Luciferase reporter assays were conducted in C2C12 myoblasts transfected with constructs containing genomic fragments harboring putative TEAD-binding motifs or with the empty pGL2-basic vector as a control. Following transfection, cells were either harvested after 48 h (myoblasts) or allowed to differentiate into myotubes for 5 days before harvesting. Cell extracts were prepared in harvest buffer (88 mM morpholineethanesulfonic acid [MES], pH 7.8; 88 mM Tris, pH 7.8; 12.5 mM magnesium acetate; 2.5 mM ATP; 1 mM DTT; 0.1% Triton X-100). Luciferase activity was measured using a Berthold CENTRO LB960 luminometer according to the manufacturer’s instructions.

### Mouse procedures and genotyping

All mouse experiments were conducted in accordance with institutional and national animal welfare regulations and were approved by the local animal ethics committees (Animal Protection Officer, Sachgebiet Tierschutzangelegenheiten, FAU Erlangen-Nürnberg; approval numbers AZ I/39/EE006, TS-07/11, and RUF-55.2.2-2532-2-1804-24) and the regional authorities (Regierung von Unterfranken). Yap1/Wwtr1 floxed mice were obtained from The Jackson Laboratory (stock no. 030532) (44). Cre-reporter mice were described previously (45,46). Mice were housed under standard conditions (22 ± 1 °C, 50-60% humidity) on a 12-h light/dark cycle with food and water ad libitum. Breeding and genotyping were performed as described earlier (47). Unless stated otherwise, adult muscles analyzed in this study were collected from mice aged 3-6 months.

### RNA-seq and expression analysis

RNA was isolated from hindlimb muscles of neonatal mice and processed using Illumina next-generation sequencing. Total RNA (500 ng per sample) was quantified and quality-checked with a 4200 Tapestation and RNA ScreenTape assay (Agilent Technologies), and libraries were prepared with the NEBNext Ultra II Directional RNA Library Preparation Kit, NEBNext Poly(A) mRNA Magnetic Isolation Module, and NEBNext Multiplex Oligos for Illumina (New England Biolabs) according to the manufacturers’ instructions. Libraries were quality-checked with a D5000 ScreenTape assay (Agilent Technologies), pooled, and sequenced on a NovaSeq 6000 SP 100-cycle v1.5 run (101-cycle, single-end, standard-loading workflow).

Base calls were converted to FASTQ format with bcl2fastq v2.20.0.422, and adapter trimming, size selection (reads > 25 nt), and quality filtering (Phred score > 43) were performed with Cutadapt. Filtered reads were aligned to the mouse genome (mm10) using Bowtie2 v2.2.9 with default settings, and read counts were obtained in R using countOverlaps (GenomicRanges). Differential expression analysis was carried out with DESeq2 v3.26.8 using default parameters, and P values were adjusted for multiple testing with the Benjamini-Hochberg procedure; genes with adjusted P < 0.05 were considered differentially expressed.

Gene Ontology (GO) enrichment analysis was performed using GOrilla (Biological Process ontology, P < 1 × 10⁻⁴), and overrepresentation analyses were carried out with WebGestaltR v0.4.6 using genes with |log₂ fold change| > 0.5 and adjusted P < 0.05.

### Statistical analysis

Statistical analyses were performed using GraphPad Prism v10 (GraphPad Software, Boston, MA, USA). Outliers were identified with Prism’s built-in outlier detection tools and excluded from further analysis as specified in the figure legends. Unless stated otherwise, comparisons between two groups were made using unpaired, two-tailed Student’s t-tests, and data are presented as mean ± SD with individual data points shown in the graphs. For comparisons involving more than two groups, one-way ANOVA followed by Dunnett’s post-hoc test was used, as detailed in the figure legends. Exact P values are reported in GraphPad format (four decimal places), and significance levels are indicated as follows: *P ≤ 0.05; **P ≤ 0.01; ***P ≤ 0.001; ****P ≤ 0.0001. Results without asterisks were considered not significant (P > 0.05).

## RESULTS

### The hindlimbs of newborn Yap1/Wwtr1 cDKO mice are shorter, whereas hindlimb bone length is preserved

Muscle-specific *Yap1*/*Wwtr1* conditional double knockout (cDKO) mice generated with HSA (human skeletal actin)-Cre have been reported to die shortly after birth, likely due to severe neuromuscular junction defects and respiratory failure (23). HSA-Cre recombinase is expressed during development shortly after myoblasts fuse to form myotubes (48). In Drosophila, loss of the Yorkie transcriptional co-activator, the homologue of YAP1, results in smaller muscles due to reduced expression of sarcomeric proteins and defective myofibril assembly (49), prompting us to examine limb growth in *Yap1*/*Wwtr1* cDKO mice. At mid-gestation (E12.5-E16.5), cDKO embryos are grossly indistinguishable from controls (Fig. 1A). By contrast, at birth the hindlimbs of cDKO mice are reduced in length by approximately 10-15% compared with control and single knockout littermates (Fig. 1B,C).

**Figure 1.**
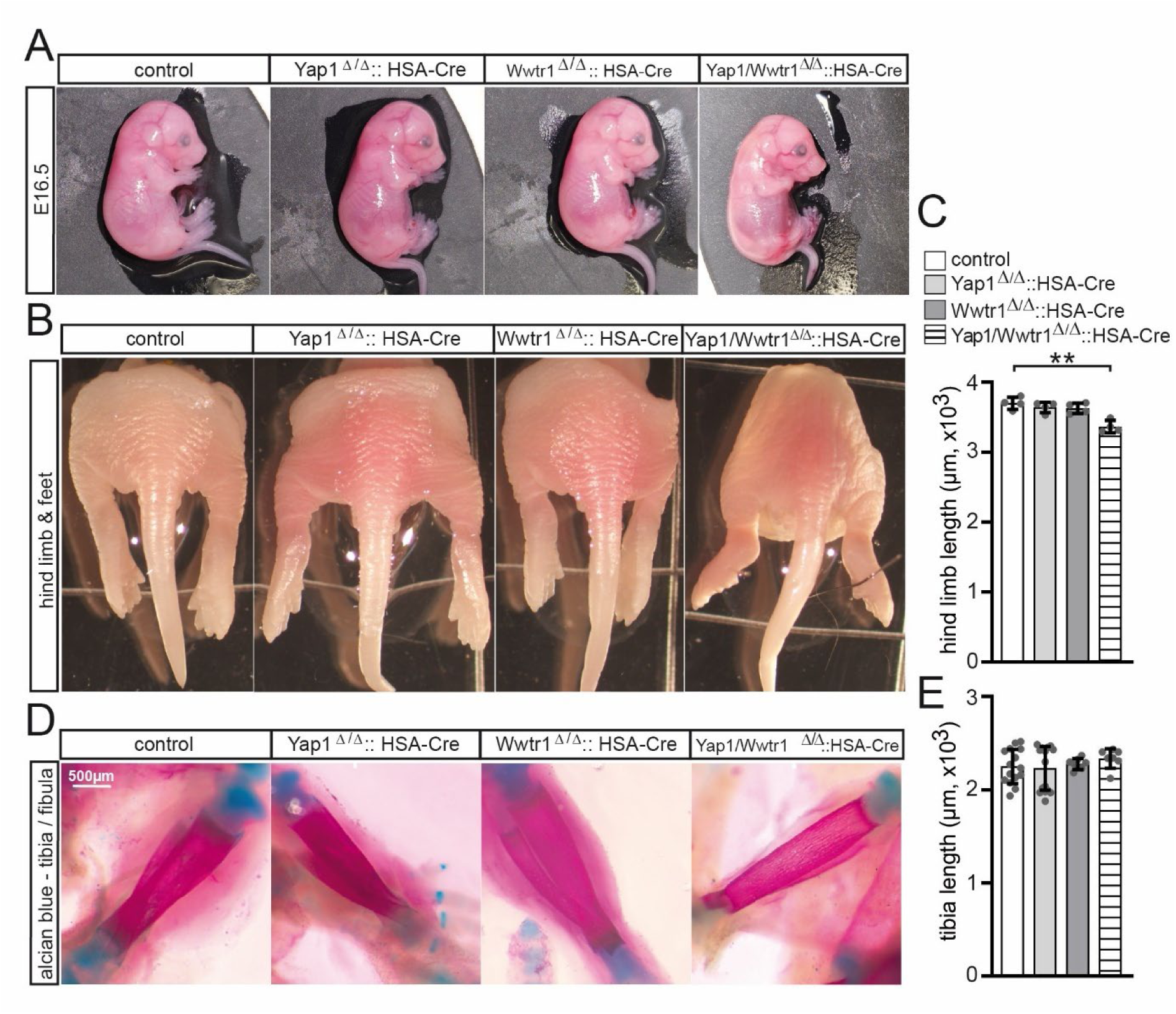
Shortened hind limbs but preserved hindlimb bone length in *Yap1*/*Wwtr1* cDKO neonates. (A) Representative images of E16.5 embryos from control, *Yap1* knockout, *Wwtr1* knockout, and *Yap1*/*Wwtr1* conditional double-knockout (cDKO) mice, showing no overt differences at this stage. (B) Hindlimb images from newborn control, single knockout, and cDKO mice. Panels show representative dorsal views of P0 hindlimbs; slight variation in skin coloration between pups reflects illumination and photographic conditions rather than overt hemorrhage or edema. (C) Quantification of hindlimb length for all genotypes, showing reduced hindlimb length in cDKO mice. (D) Representative skeletal preparations of newborn tibiae from all genotypes stained with Alcian blue. (E) Quantification of tibia length for all genotypes, demonstrating that tibia length is preserved despite reduced overall hindlimb size in cDKO mice. Data in (C,E) are mean ± SD (n ≥ 3 mice per genotype). Statistical significance was assessed using unpaired, two-tailed Student’s t-tests.

Because overall limb length is primarily determined by long bone growth, whereas apparent shortening can also arise from tendon retraction secondary to muscle loss, we next quantified hindlimb bone length. Alcian blue staining of tibia and fibula preparations from newborn control, single knockout, and cDKO mice revealed normal skeletal patterning and unchanged tibia length in cDKO animals (Fig. 1D,E). These findings indicate that the shortened hindlimbs of *Yap1*/*Wwtr1* cDKO mice are not due to intrinsic defects in long bone growth, but are most likely secondary to impaired skeletal muscle development.

### Embryonic MYH staining reveals abnormal hindlimb muscle development and reduced muscle size in Yap1/Wwtr1 cDKO mice

We next asked whether delayed muscle development and/or growth contributes to the reduced hindlimb length in *Yap1*/*Wwtr1* cDKO mice. To this end, we analyzed cross-sections of hindlimbs from control, single knockout, and cDKO embryos at E14.5, E16.5, E18.5, and P0, and quantified the cross-sectional area of individual muscles (SOL, soleus; GAS, gastrocnemius; EDL, extensor digitorum longus; TA, tibialis anterior) on sections with maximal muscle size (Supplementary Fig. 1, Fig. 2A). MYH3 immunostaining, which broadly labels embryonic myofibers and remains detectable around birth, was used to visualize the overall muscle architecture, while MYH7 staining (labels slow type fibers) at E14.5 highlighted fibers derived from the first wave of myogenesis (Supplementary Fig. 1).

**Figure 2.**
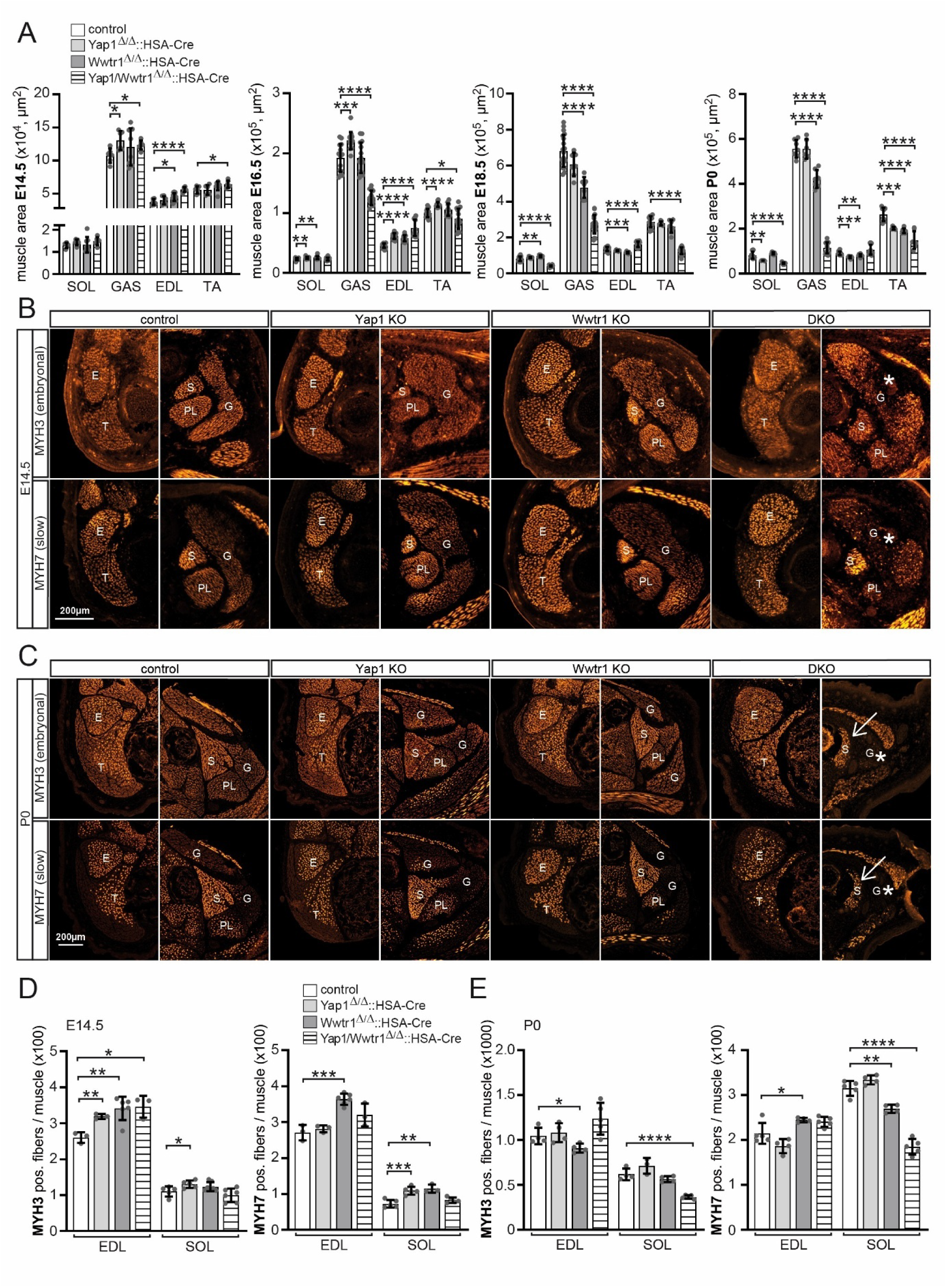
*Yap1*/*Wwtr1* deficiency reduces hindlimb muscle size and impairs embryonic-early postnatal fiber differentiation. (A) Quantification of muscle cross-sectional area (SOL, GAS, EDL, and TA) at E14.5, E16.5, E18.5, and P0 in control, single knockout, and double-knockout mice. Muscle area is reduced in cDKO muscles from E16.5 onwards. (B) Representative transverse hindlimb sections stained for MYH3 and MYH7 at E14.5 from all genotypes, showing impaired GAS development in cDKO embryos. Major hindlimb muscles, including soleus (S), plantaris (PL), extensor digitorum longus (E), gastrocnemius (G), and tibialis anterior (T), are indicated. Asterisks highlight absence or marked reduction of the medial GAS. Scale bar, 200 µm. (C) Representative transverse hindlimb sections stained for MYH3 and MYH7 at P0, showing persistent muscle size reduction in cDKO muscles. Arrows indicate the smaller and morphologically altered SOL; asterisks indicate impaired GAS development. Muscle groups are labeled as in (B). Scale bar, 200 µm. (D) Quantification of MYH3- and MYH7-positive fibers per muscle at E14.5 in EDL and SOL, showing increased numbers of early myosin isoform-positive fibers in Yap1-deficient, Wwtr1-deficient, and cDKO embryos. (E) Quantification of MYH3- and MYH7-positive fibers per muscle at P0 in EDL and SOL. cDKO muscles show the strongest reduction in fiber number in the SOL, indicating impaired early fiber differentiation. Analysis and labeling as in (D). Data are mean ± SD (n ≥ 3 mice per genotype). Statistical significance was assessed using unpaired, two-tailed Student’s t-tests.

At E14.5, MYH3 staining revealed similar cross-sectional areas for most hindlimb muscles in control and single knockout embryos, whereas cDKO hindlimbs already showed impairment of the central region of the GAS, which later forms the medial head (Supplementary Fig. 1 labeled by asteriks, Fig. 2A). MYH7 staining and muscle positioning were comparable between control, *Yap1*-deficient, and *Wwtr1*-deficient embryos; in contrast, cDKO hindlimbs lacked MYH7-positive fibers in the GAS and plantaris, whereas other muscles were only mildly or not appreciably affected. (Supplementary Fig. 1).

By E16.5, MYH3 staining indicated that muscle regions remained largely intact in control and single knockout embryos, and separation of the GAS into medial and lateral heads was clearly visible. In cDKO hindlimbs, this separation was never observed; the GAS was markedly reduced and structurally disrupted, with MYH3-negative regions within the expected GAS muscle area (Supplementary Fig. 1, Fig. 2A). From E16.5 onwards, the plantaris was no longer detectable in cDKO embryos, and the deep flexor and peroneal muscle groups were reduced, with some constituent muscles missing. MYH7 staining in control and single knockouts showed enrichment of slow fibers toward the bone and more peripheral fast-fiber regions, consistent with the second wave of myogenesis, whereas this pattern was only partially retained in cDKO EDL and TA and largely absent in other muscles. MYH7-positive fibers were essentially absent in the GAS and markedly reduced in the TA, deep flexor, and peroneal muscles of cDKO hindlimbs (Supplementary Fig. 1).

At E18.5, control and single knockout embryos displayed characteristic MYH3 patterns, with stronger staining in TA and EDL regions adjacent to the bone and a more uniform distribution in the SOL (Supplementary Fig. 1). In cDKO hindlimbs, MYH3-negative or weakly stained regions in TA and EDL were largely lost, and several muscles were either reduced or absent, leading to altered spatial relationships among the remaining muscles (Supplementary Fig. 1, Fig. 2A). For example, flexor digitorum longus shifted position, likely due to loss of deep flexor muscles, and the SOL was reduced and deformed, consistent with the absence of surrounding GAS and plantaris; peroneal muscles also began to vanish in cDKO embryos. MYH7-positive fibers were nearly absent in TA, undetectable in GAS, and significantly reduced in multiple other cDKO muscles (Supplementary Fig. 1).

At birth, MYH3 staining showed preserved overall muscle patterning in single knockout hindlimbs, although both *Yap1* and *Wwtr1* single knockouts exhibited reduced GAS and/or TA areas (Supplementary Fig. 1, Fig. 2A). In cDKO hindlimbs, muscle defects were region-specific: EDL area was largely maintained, whereas GAS, plantaris, deep flexor, and peroneal muscles were almost undetectable by MYH3 staining, and flexor digitorum longus was displaced and reduced (Supplementary Fig. 1, Fig. 2A). The SOL was markedly smaller and altered in shape, likely secondary to the loss of adjacent muscles. MYH7-positive fibers remained detectable in EDL and SOL but were strongly reduced in most other residual muscles.

To quantify how these architectural defects relate to fiber formation, we determined the number of MYH3- and MYH7-positive fibers in EDL and SOL at E14.5 and at birth (Fig. 2B-E). We focused on these muscles because EDL is predominantly fast-twitch and rapidly transitions from embryonic MYH3 to adult fast myosins (MYH1/2/4), whereas SOL becomes enriched in slow MYH7, allowing us to assess maturation of embryonic MYH3 fibers into slow MYH7 fibers in a slow-biased environment. At E14.5, the number of MYH3-positive fibers in EDL was slightly increased in both single knockouts and cDKO compared with controls, whereas SOL fiber counts were largely unchanged (Fig. 2B,D). MYH7-positive fiber numbers showed a modest increase in *Wwtr1*-deficient EDL and SOL (Fig. 2B,D).

At P0, MYH3-positive fiber numbers in EDL were similar or slightly reduced in *Wwtr1* knockouts relative to controls, whereas SOL muscles of cDKO neonates showed a pronounced reduction in MYH3-positive fibers, consistent with the marked decrease in SOL area (Fig. 2A,C,E). A similar trend was observed for MYH7: SOL from *Wwtr1*-deficient and cDKO mice showed reduced numbers of MYH7-positive fibers, whereas EDL from *Wwtr1* knockouts exhibited a modest increase (Fig. 2C,E).

Taken together, these observations indicate that initial formation of the first wave of myofibers is largely preserved in *Yap1*/*Wwtr1* cDKO hindlimbs at E14.5, but subsequent growth and maturation of multiple muscle groups are severely compromised. As development proceeds, this culminates in the near-complete loss of several distal muscles such as the gastrocnemius and plantaris and a marked reduction in slow fiber formation in the SOL.

### YAP1 and TAZ drive myotube formation and fiber growth across primary and secondary waves of myogenesis

During embryonic myogenesis, primary myotubes form first and rapidly enlarge, whereas secondary myotubes form later, initially as small fibers attached to the primary scaffolds, and progressively grow until the size distinction between the two waves is lost. To pinpoint which steps of this process require YAP1 and TAZ, we analyzed individual MYH3-stained fibers on cross-sections of EDL and SOL muscles from control, single knockout, and *Yap1*/*Wwtr1* cDKO embryos at E14.5, E16.5, and E18.5 (Fig. 3).

**Figure 3.**
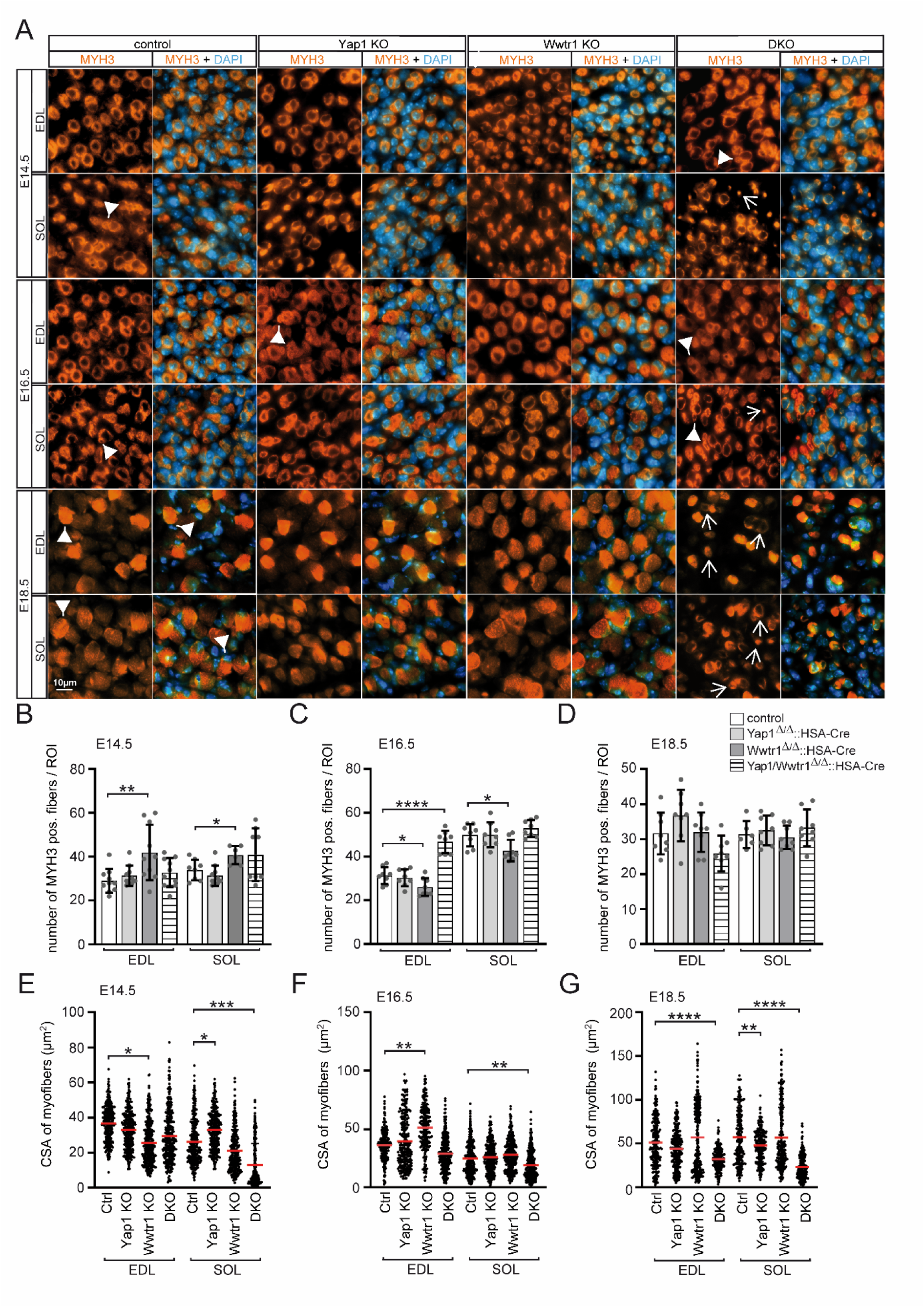
YAP1 and TAZ drive myotube formation and fiber growth across primary and secondary waves of myogenesis. (A) Representative high-magnification images of MYH3-stained myofibers (orange) with DAPI-stained nuclei (blue) in EDL and SOL muscles from control, single knockout, and cDKO embryos at E14.5, E16.5, and E18.5. White arrowheads indicate normally maturing primary and secondary myofibers with peripheral nuclei and proper attachment. In cDKO muscles, arrows highlight impaired fiber maturation, including reduced cross-sectional area (CSA), failure of secondary fiber attachment, and persistent doughnut-shaped morphology. Scale bar, 10 µm. (B-D) Quantification of MYH3-positive fibers per region of interest (ROI) in EDL and SOL at E14.5 (B), E16.5 (C), and E18.5 (D). Note the transient increase in fiber number in Wwtr1-deficient and cDKO muscles at early stages, followed by a decline at E16.5 and E18.5. Data are presented as mean ± SD (n ≥ 4 mice per genotype). Figure legend as shown in (D). Statistical significance was assessed using an unpaired Student’s t-test. (E-G) Distribution of myofiber CSA in EDL and SOL at E14.5 (E), E16.5 (F), and E18.5 (G) shown as violin plots. Each dot represents the individual MYH3-positive fiber CSA per muscle; statistical analysis was performed on the mean fiber size per mouse (n ≥ 4 per genotype). CSA distributions in single knockouts remain largely comparable to controls, whereas cDKO fibers show a marked shift toward smaller CSAs across all stages, most prominently in the SOL. Red horizontal lines indicate median values. Statistical significance was assessed by one-way ANOVA followed by Dunnett’s post-hoc test.

At E14.5, EDL from control and *Yap1* knockout embryos contained comparable numbers of large, doughnut-shaped primary fibers, although MYH3-positive fibers in Yap1 mutants showed a modestly reduced cross-sectional area (CSA; Fig. 3A,B,E). *Wwtr1* knockout EDL displayed fewer large-diameter fibers but more small secondary fibers clustered around primary fibers, resulting in an increased total fiber count per region of interest (ROI). Similarly, cDKO EDL exhibited elevated fiber numbers with smaller CSA, and many small fibers appeared attached to large doughnut-shaped primaries (Fig. 3A, white arrowhead). In SOL at E14.5, control embryos showed large primary myotubes surrounded by smaller secondary fibers (depicted by white arrowhead). *Yap1* knockout SOL contained slightly fewer MYH3-positive fibers but with increased CSA, while secondary fibers remained appropriately associated with primary scaffolds (Fig. 3A,B,E). *Wwtr1*-deficient SOL resembled Wwtr1-deficient EDL, with increased clustering of secondary fibers around primary myotubes. In cDKO SOL, MYH3-positive fiber number was similar to control but CSA was markedly reduced; notably, some small nascent fibers appeared randomly distributed and not attached to primary scaffolds, a defect less pronounced in EDL (Fig. 3A, white arrow).

At E16.5, EDL from *Yap1* knockouts closely resembled controls in both MYH3-positive fiber number and CSA, and secondary fibers remained clustered around large doughnut-shaped primaries (labeled by white arrowheads, Fig. 3A,C,F). In contrast, *Wwtr1* knockout EDL showed fewer MYH3-positive fibers per ROI but an increased mean CSA, together with a reduction in small secondary fibers attached to primary scaffolds (Fig. 3A,C,F). cDKO embryos displayed a transient increase in MYH3-positive fiber density, but these fibers were predominantly small, leading to a heterogeneous population with reduced CSA. Many secondary fibers still clustered around the remaining large primary myotubes (white arrowhead), but in SOL a substantial fraction of small fibers failed to attach to primary scaffolds and appeared randomly positioned (Fig. 3A, arrow), while others remained attached (Fig. 3A, arrowheads). Thus, at E16.5, SOL most clearly illustrates the failure of secondary fibers to integrate into the primary scaffold network in the absence of YAP1/TAZ, whereas *Wwtr1* single loss mainly shifts the balance toward fewer but larger fibers.

By E18.5, large doughnut-shaped primary myotubes in control, *Yap1*, and *Wwtr1* knockout EDL had largely matured, appearing fully filled with MYH3 signal and exhibiting peripheral nuclei (white arrowhead), although small nascent secondary fibers with central nuclei remained detectable (Fig. 3A). Total numbers of MYH3-positive fibers in EDL and SOL were similar between single knockouts and controls (Fig. 3D), but *Wwtr1*-deficient EDL contained a highly heterogeneous fiber population with clusters of very small secondary fibers surrounding abnormally large primaries. In cDKO muscles, MYH3-positive fiber numbers decreased relative to E16.5, and nearly all remaining fibers, including secondary fibers, displayed reduced CSA (Fig. 3G). This drop in fiber number between E16.5 and E18.5 suggests a progressive loss of immature myofibers that fail to mature. Although many primary myotubes in cDKO EDL eventually filled with MYH3 signal, persistent doughnut-shaped fibers with MYH3-negative centers were still observed (Fig. 3A, white arrow). The phenotype was even more striking in cDKO SOL: total fiber number remained roughly constant, but CSA was severely reduced and most fibers retained a doughnut-like morphology with empty centers and high heterogeneity (white arrow, Fig. 3A,D,G).

Given the perinatal lethality of the cDKO, we also examined adult single knockout muscles to assess long-term consequences of *Yap1* or *Wwtr1* loss. While single knockouts were viable, both showed distinct structural changes (Supplementary Fig. 2). Cytochrome oxidase staining did not indicate a major metabolic shift (Supplementary Fig. 2A), but *Wwtr1* mutants displayed a pronounced fiber-type shift, with an increased proportion of MYH7-positive type I fibers and fewer MYH2-positive type IIa fibers (Supplementary Fig. 2B,D). Both single knockouts showed elevated percentages of centrally located nuclei, more prominently in *Wwtr1* mutants (Supplementary Fig. 2A,C), and opposing effects on fiber size, *Yap1* deficiency reduced CSA across fiber types, whereas *Wwtr1* loss led to larger fibers (Supplementary Fig. 2A,E). These adult phenotypes parallel the embryonic observations and support non-redundant roles for YAP1 and TAZ in regulating fiber growth and fiber-type composition, with TAZ playing a particularly prominent role in slow muscle. A detailed dissection of TGFβ signaling and of protein synthesis and degradation pathways in these single knockout muscles is beyond the scope of the present study and will require dedicated future work.

### Global myogenic transcript changes in Yap1/Wwtr1 double-knockout muscle identify candidate YAP1/TAZ-TEAD target genes

Because YAP1 and TAZ act as transcriptional co-regulators, we performed RNA-sequencing on hindlimb muscles from newborn control, *Yap1*-deficient, *Wwtr1*-deficient, and *Yap1*/*Wwtr1* cDKO mice to identify YAP/TAZ-dependent gene expression changes (Fig. 4). Principal component analysis revealed that transcriptomes from single knockouts clustered closely with controls, whereas cDKO samples formed a clearly separated group, indicating a distinct transcriptional state (Fig. 4A). Consistently, *Yap1*-deficient muscles showed no upregulated genes and only four downregulated genes, and *Wwtr1* deficiency led to 26 upregulated and 70 downregulated genes (padj < 0.05), while the cDKO caused extensive dysregulation with 543 upregulated and 1,539 downregulated genes (Fig. 4B,C).

**Figure 4.**
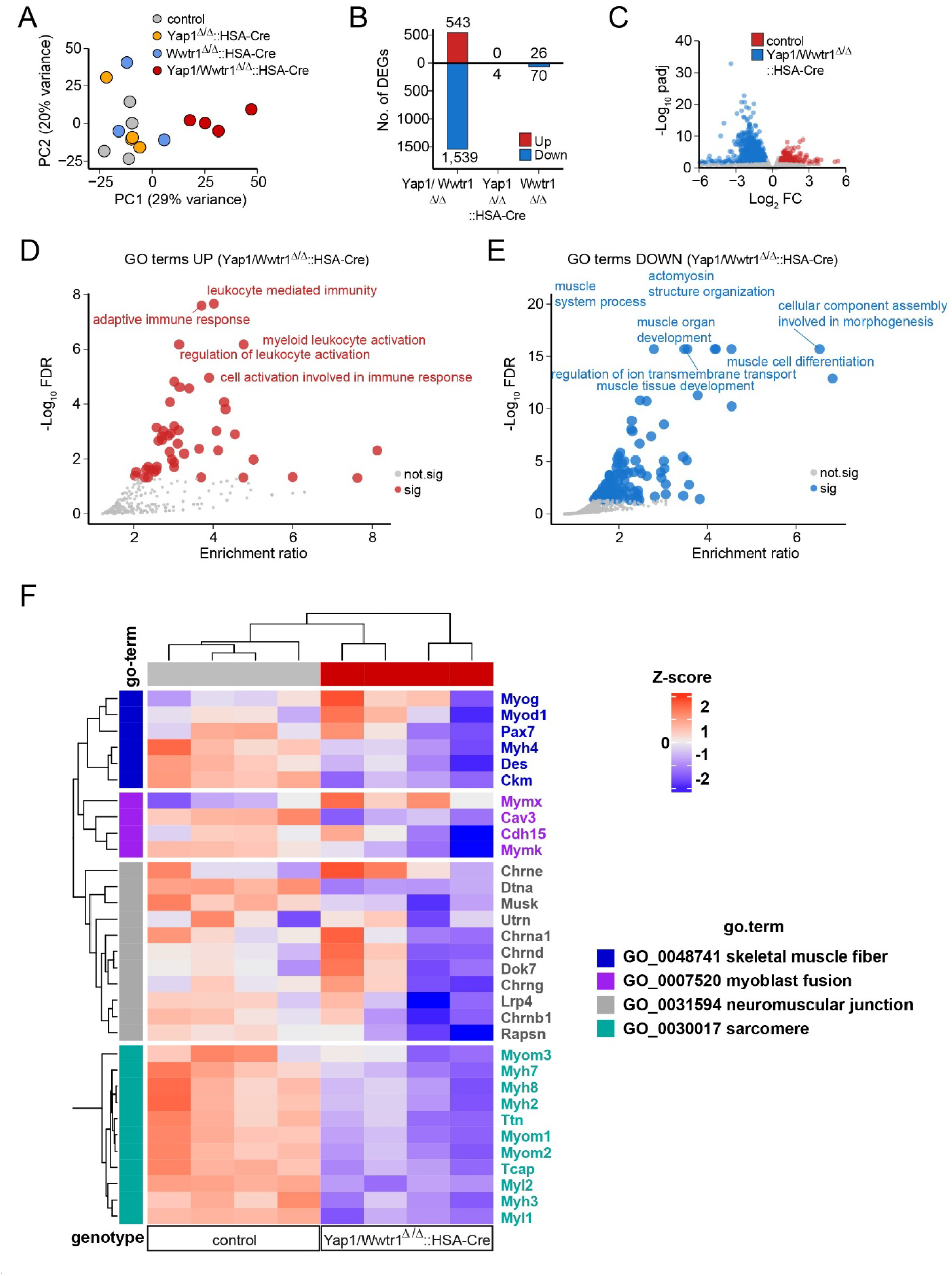
RNA-seq analysis of Yap1/Wwtr1 double-knockout soleus muscle identifies myogenic YAP1/TAZ-TEAD-dependent target genes. (A) Principal component analysis of transcriptomes from the indicated genotypes (n = 4 per group). (B) Number of differentially expressed genes (DEGs; padj < 0.05) per genotype. Single knockouts show only limited transcriptional changes, whereas the cDKO exhibits 543 upregulated and 1,539 downregulated genes. (C) Volcano plot showing transcripts significantly up- or downregulated in cDKO versus control muscle. Log2FC, log2 fold change; padj, adjusted P value. (D) Gene Ontology (GO) terms enriched among upregulated genes in the cDKO, including immune response-related categories such as leukocyte-mediated immunity and myeloid leukocyte activation. (E) GO terms enriched among downregulated genes, including muscle fiber development, muscle structure organization, and cellular component assembly involved in morphogenesis. (F) Heatmap of expression profiles for key muscle markers and myogenesis-associated genes. Red indicates high expression; blue indicates low expression. Gene sets correspond to skeletal muscle fiber, myoblast fusion, neuromuscular junction, and sarcomere-associated categories.

Gene Ontology analysis of cDKO-specific DEGs showed that upregulated genes were strongly enriched for immune-related categories, including leukocyte-mediated immunity and myeloid leukocyte activation, consistent with a secondary inflammatory response to muscle pathology (Fig. 4D). In contrast, downregulated genes clustered in GO terms linked to muscle development, muscle fiber differentiation, actomyosin structure organization, and cellular component assembly involved in morphogenesis (Fig. 4E). To focus on processes most relevant to our phenotypes, we generated a heatmap of representative genes associated with “skeletal muscle fiber,” “myoblast fusion,” “neuromuscular junction,” and “sarcomere” categories (Fig. 4F). This analysis highlights a broad reduction in transcripts for sarcomeric and postsynaptic genes, as well as selected myogenic regulators.

In particular, nearly all myosin heavy chain genes, which encode thick-filament components, were strongly downregulated in cDKO muscles. Transcripts for myosin light chains, titin (Ttn), and multiple myomesin and palladin family members were also reduced. Among fusion-related genes, Myomaker (*Mymk*) and several other myoblast fusion factors were markedly decreased, correlating with the impaired myotube formation seen *in vivo*. Several genes involved in muscle structure and function, such as Myh4, desmin (*Des*), muscle creatine kinase (*Ckm*), and caveolin-3 (*Cav3*), and key postsynaptic genes at the neuromuscular junction, were likewise downregulated explaining previously reported absence of NMJs in cDKO mice (Fig. 4F) (23).

Together, these data show that combined loss of *Yap1* and *Wwtr1* triggers a broad transcriptional collapse of myogenic fusion and structural programs, whereas single-gene deletions have relatively modest effects at birth. This transcriptomic signature provided the basis for our subsequent integration with TEAD1/4 binding and chromatin accessibility data to identify likely YAP1/TAZ-TEAD target genes involved in myotube formation and sarcomere assembly. Because cDKO muscles are smaller and contain fewer nuclei, we sequenced equal RNA amounts from each sample. Consequently, our RNA-seq analysis reflects relative expression changes per unit of remaining muscle tissue, rather than absolute transcript numbers per whole muscle.

### Combined loss of Yap1 and Wwtr1 impairs myotube formation and disrupts sarcomere architecture

The developmental abnormalities in several hindlimb muscles suggested that myotube formation may be compromised in *Yap1*/*Wwtr1* cDKO mice. Because *Mymk* (Myomaker), a key fusion gene, was strongly downregulated in cDKO muscles (Fig. 4F), we tested whether YAP1/TAZ are required cell-intrinsically for myotube formation. Primary myogenic cells were isolated from Pax7-CreER^T2^ mice, and *Yap1*/*Wwtr1* deletion was induced *ex vivo* by tamoxifen treatment. Upon differentiation, control cultures formed numerous elongated, multinucleated myotubes, whereas tamoxifen-induced *Yap1*/*Wwtr1*-deficient cultures contained only sparse myotubes despite a comparable density of mononuclear cells (Fig. 5A). Quantification confirmed a marked reduction in the number of myotubes per field (Fig. 5B) and showed that myonuclear incorporation per myotube remained significantly lower than in controls (Fig. 5C), indicating a cell-intrinsic fusion defect in the absence of YAP1/TAZ.

**Figure 5.**
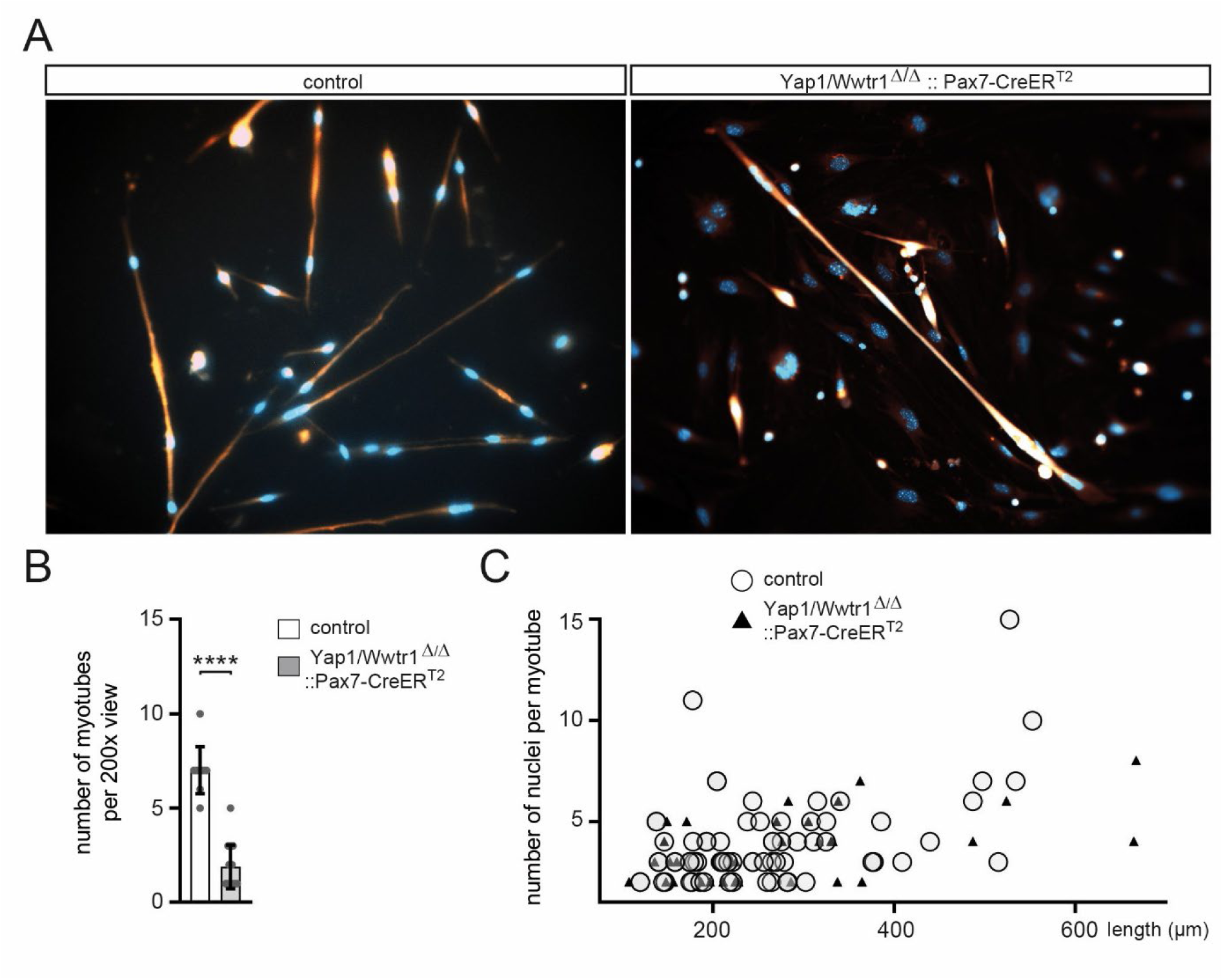
Loss of YAP1/TAZ impairs myotube formation *ex vivo*. (A) Representative immunofluorescence images of differentiated primary myoblast cultures derived from control or *Yap1*/*Wwtr1^Δ^*^/Δ^::Pax7-CreER^T2^ mice, showing myotubes (BTX, red) and nuclei (DAPI, blue) after induction of differentiation. (B) Quantification of the number of myotubes per 200× field in control and *Yap1*/*Wwtr1^Δ^*^/Δ^::Pax7-CreER^T2^ cultures (mean ± SD; each dot represents one field from an independent culture). Statistical significance was assessed using an unpaired Student’s t-test. (C) Scatter plot of myotube length versus the number of nuclei per myotube for control (open circles) and *Yap1*/*Wwtr1^Δ^*^/Δ^::Pax7-CreER^T2^ (black triangles) cultures, illustrating reduced myonuclear accretion in *Yap1*/*Wwtr1*-deficient myotubes.

In addition to impaired fusion, our transcriptomic data from cDKO SOL muscles revealed broad downregulation of sarcomere-associated genes, consistent with previous reports that loss of the YAP1 orthologue Yorkie disrupts sarcomere organization in Drosophila flight muscle (49). We therefore re-examined sarcomere structure in neonatal diaphragm at higher resolution. Light microscopy of P0 diaphragm showed regularly aligned myofibrils with well-defined striations in control and single knockout fibers, whereas *Yap1*/*Wwtr1* cDKO fibers displayed fewer discernible sarcomeres and irregular, patchy striation patterns (white arrows, Fig. 6A).

**Figure 6.**
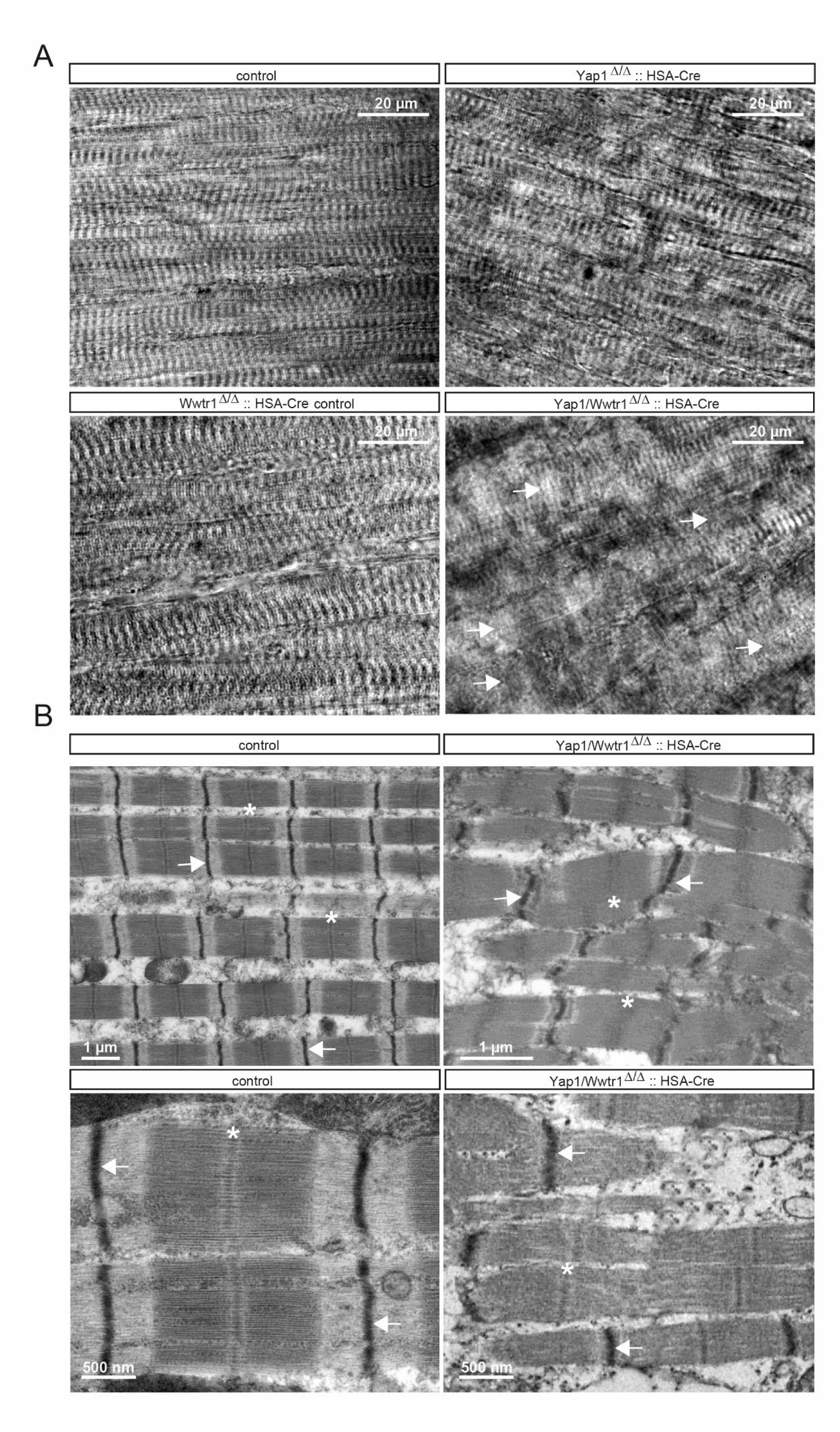
Hippo pathway inactivation disrupts sarcomere organization and triad ultrastructure in skeletal muscle. (A) Light microscopy of P0 diaphragm shows regularly aligned myofibrils with well-defined striations in control and single knockout fibers, whereas Yap1/Wwtr1 cDKO fibers display fewer discernible sarcomeres and irregular, patchy striation patterns (arrow). Representative images from n ≥ 3 mice per genotype. (B) Transmission electron microscopy of newborn diaphragm muscle reveals intact sarcomeres with sharply defined Z-lines (arrow), A-bands (asterisk), and I-bands in control fibers and disorganized sarcomeres with broadened Z-lines (arrow), misaligned A-bands (asterisk), and areas of myofibrillar disassembly in cDKO fibers. Representative images from n ≥ 3 mice per genotype. Scale bars: 20 µm (A); 2 µm (top), 500nm (bottom) (B).

Transmission electron microscopy further highlighted these defects at the ultrastructural level. Control fibers exhibited intact sarcomeres with sharply defined Z-lines, A-bands, and I-bands, whereas cDKO fibers showed broadened and distorted Z-line regions, misaligned A-bands, and areas of myofibrillar disassembly (Fig. 6B). These abnormalities indicate that YAP1/TAZ activity in differentiated myofibers is required not only for proper sarcomere maturation but also for maintenance of myofibrillar integrity *in vivo*.

Collectively, the *ex vivo* fusion assays and *in vivo* structural analyses demonstrate that combined loss of *Yap1* and *Wwtr1* compromises both efficient myotube formation and sarcomere organization, supporting a dual role for YAP1/TAZ in building and stabilizing the contractile apparatus of skeletal muscle fibers.

### Integrating Yap1/Wwtr1 double-knockout transcriptomes with ECR and TEAD ChIP-seq data reveals TEAD-linked myogenic regulation

To assess whether genes dysregulated upon *Yap1*/*Wwtr1* loss are associated with TEAD regulatory activity during myogenesis, we compared the cDKO neonatal muscle transcriptome with published TEAD1 and TEAD4 ChIP-seq datasets from C2C12 cells at day 0 and day 6 of differentiation (34). Genes harboring TEAD1 or TEAD4 peaks within 1 kb or 50 kb of their transcription start sites showed a clear shift toward reduced expression in cDKO muscle, whereas genes without nearby TEAD binding sites did not exhibit a comparable shift in log2 fold change (Fig. 7A). This pattern was observed for both TEAD1 and TEAD4 at day 0 and day 6, suggesting that TEAD-proximal genes are disproportionately sensitive to the loss of YAP1/TAZ activity and likely belong to transcriptional programs positively regulated by the YAP1/TAZ-TEAD network.

**Figure 7.**
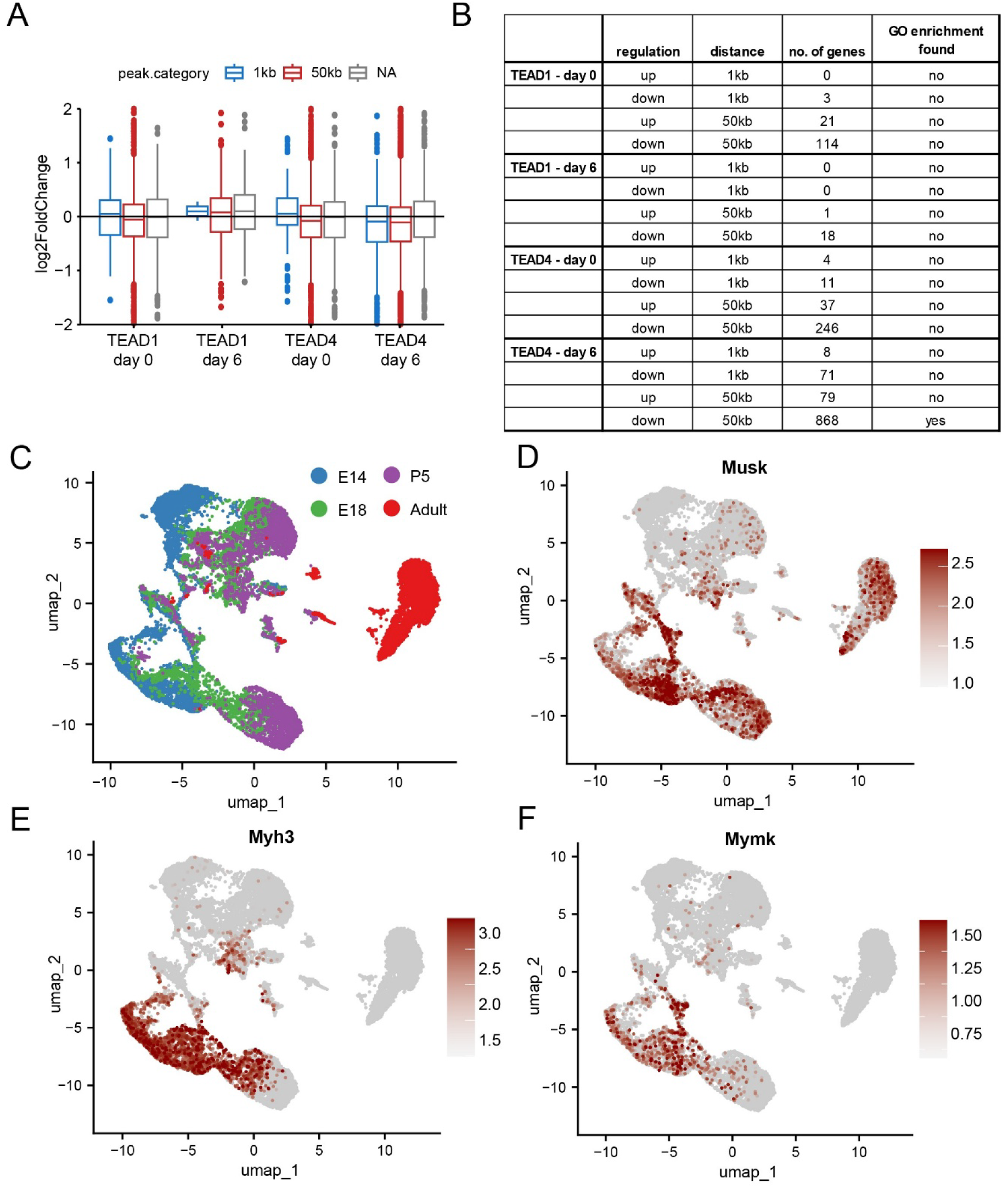
Temporal analysis of TEAD binding and gene expression during muscle development. (A) Box plots showing the log2 fold-change in gene expression in cDKO muscles for genes linked to M-CAT-containing TEAD1/4 binding sites reported by ChIP-seq (34). Genes were grouped according to the distance of the nearest TEAD peak from the transcription start site (1 kb, 50 kb, or NA), and TEAD1 or TEAD4 binding is shown at day 0 and day 6 of C2C12 differentiation. (B) Summary table of differential gene regulation, number of genes, and GO term enrichment for genes associated with TEAD1/4 peaks within 1 kb or 50 kb of the transcription start site at the indicated time points (TEAD1 day 0, TEAD1 day 6, TEAD4 day 0, TEAD4 day 6). (C) UMAP plot showing clustering of muscle nuclei at different developmental stages—E14 (embryonic day 14), E18 (embryonic day 18), P5 (postnatal day 5), and adult—modified from (35). Clusters are color-coded by developmental stage. (D-F) UMAP plots showing the expression patterns of Musk (D), Myh3 (E), and Mymk (F) across the clusters. Expression levels are indicated by the color scale, with higher expression shown in darker red.

Quantitative analysis of TEAD-bound genes showed that relatively few upregulated genes overlapped TEAD1/4 binding sites, whereas a large fraction of downregulated genes - especially those with TEAD4 peaks within 50 kb - overlapped these sites. (Fig. 7B). GO enrichment was detected specifically among downregulated genes linked to TEAD4 binding sites at day 6, emphasizing a functional connection between TEAD4 occupancy and differentiation-related pathways.

To place these TEAD-linked genes within the cellular trajectory of myogenesis, we examined their expression across a developmental single-cell RNA-seq atlas of mouse skeletal muscle (35). UMAP projection of embryonic (E14, E18), early postnatal (P5), and adult muscle nuclei recapitulated distinct developmental and mature myogenic compartments (Fig. 7C). Representative TEAD-bound genes with markedly altered transcript levels, including Musk, Myh3, and Mymk, showed restricted, stage-appropriate expression in differentiating and terminally fusing myocytes (Fig. 7D-F). Musk was enriched in neuromuscular junction-associated populations, whereas *Myh3* and *Mymk* were expressed in late embryonic and postnatal myofibers undergoing growth and maturation. The strong down-regulation of these genes in cDKO muscle, together with their localization along the myogenic trajectory, supports a model in which YAP1/TAZ-TEAD activity sustains expression of key fusion and structural genes required for terminal differentiation.

### Conserved TEAD-regulatory motifs control expression of genes involved in myotube differentiation and sarcomere structure

TEAD transcription factors regulate gene expression by binding M-CAT DNA motifs, which are enriched in regulatory regions of many muscle-specific genes and have been implicated in control of myogenic and synaptic programs (50–52). A recent study profiles TEAD function in myogenesis generated ChIP-seq datasets for TEAD1 and TEAD4 in undifferentiated and differentiated C2C12 cells (34). We re-analyzed these datasets to assess TEAD1/4 occupancy at promoter and enhancer regions of myogenic genes, extending our previous approach used to identify postsynaptic TEAD targets (23). This analysis revealed strong TEAD4 enrichment at several key myogenic loci—including the fusion gene *Myomaker* and multiple sarcomere genes encoding myosin light and heavy chains and titin (Supplementary Table 2). Several of these candidates were selected for further functional interrogation based on their essential roles in muscle differentiation and contractile architecture. Consistent with our earlier work on synaptic genes, TEAD4, but not TEAD1, occupied these regulatory regions despite the high similarity of their DNA-binding motifs (34).

We next used the JASPAR 2018 database to search these TEAD4-occupied intervals for evolutionarily conserved TEAD-binding sites across mouse, rat, dog, and human genomes (42). Representative motifs are shown in Figures 8 and 9, with a complete list in Supplementary Table 3. Integration with available single-cell ATAC-seq datasets from human skeletal muscle demonstrated that these predicted TEAD motifs reside within open chromatin regions in myocytes (Figs. 8, 9) (53), supporting their potential function as active regulatory elements *in vivo*.

**Figure 8.**
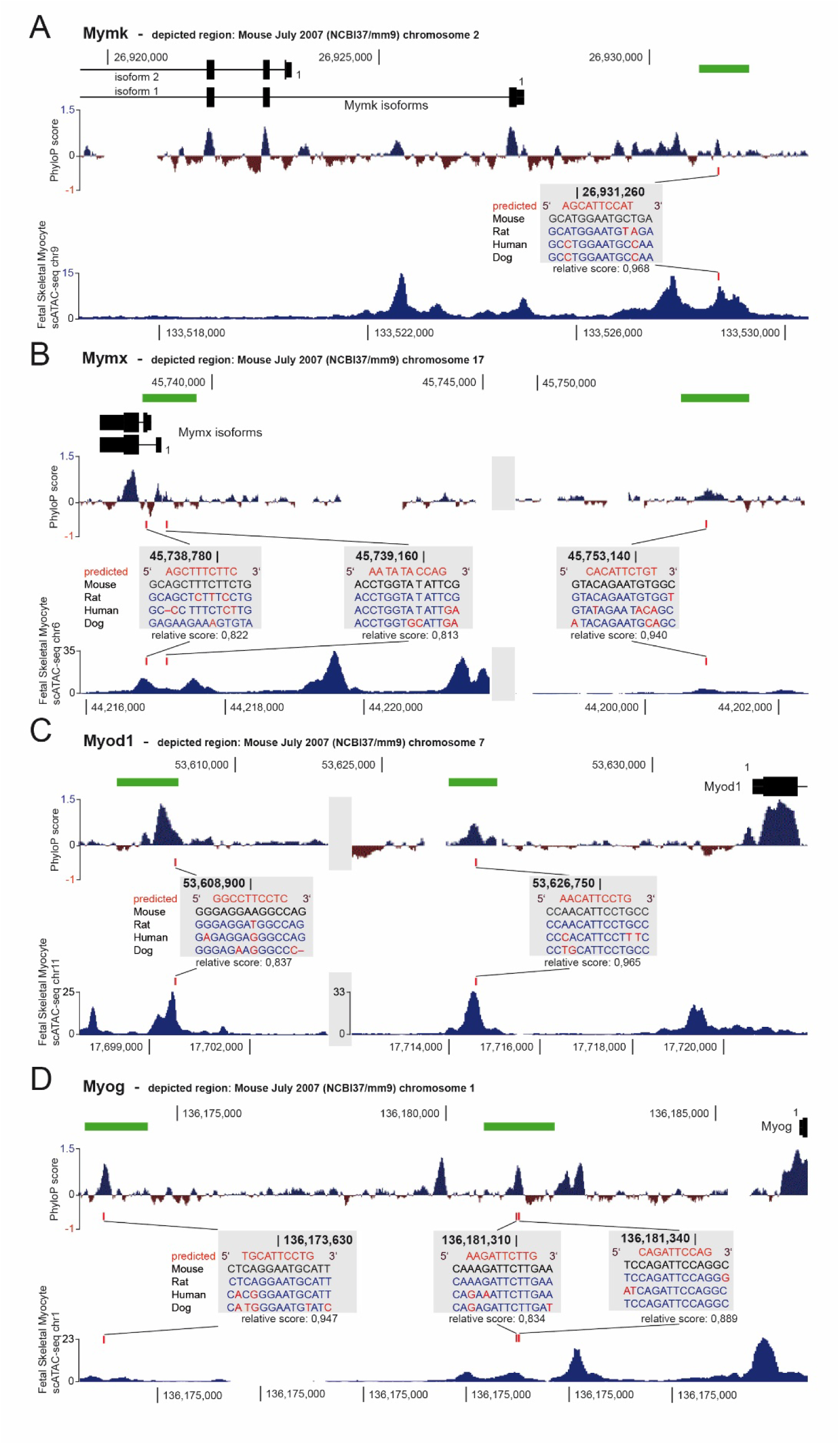
TEAD-bound M-CAT motifs in conserved regions of myogenic genes Mymk, Mymx, Myod1, and Myog. TEAD binding sites regulate expression of Mymk, Mymx, Myod1, and Myog by engaging M-CAT motifs in evolutionarily conserved regulatory regions of these myogenic genes. Putative M-CAT motifs for TEAD transcription factors were searched within evolutionarily conserved regions that are occupied by TEAD4 in ChIP-seq experiments from differentiated C2C12 myotubes. Several highly conserved candidate M-CAT sites were identified in Mymk (A), Mymx (B), Myod1 (C), and Myog (D). Each genomic locus is shown with PhyloP conservation scores, highlighting exons and potential regulatory regions. TEAD4-occupied sites are indicated as bright green rectangles, whereas predicted TEAD4 binding sites are marked in red and shown in boxes with multi-species sequence alignments. Putative TEAD4 sites were identified using the JASPAR 2018 Scan function (TEAD4 matrix profile ID MA0809.1) with a relative score threshold of 80%; the relative score for each site is indicated below the alignment. Single-cell ATAC-seq data further demonstrate that these M-CAT motifs reside within open chromatin regions in human skeletal myocytes.

**Figure 9.**
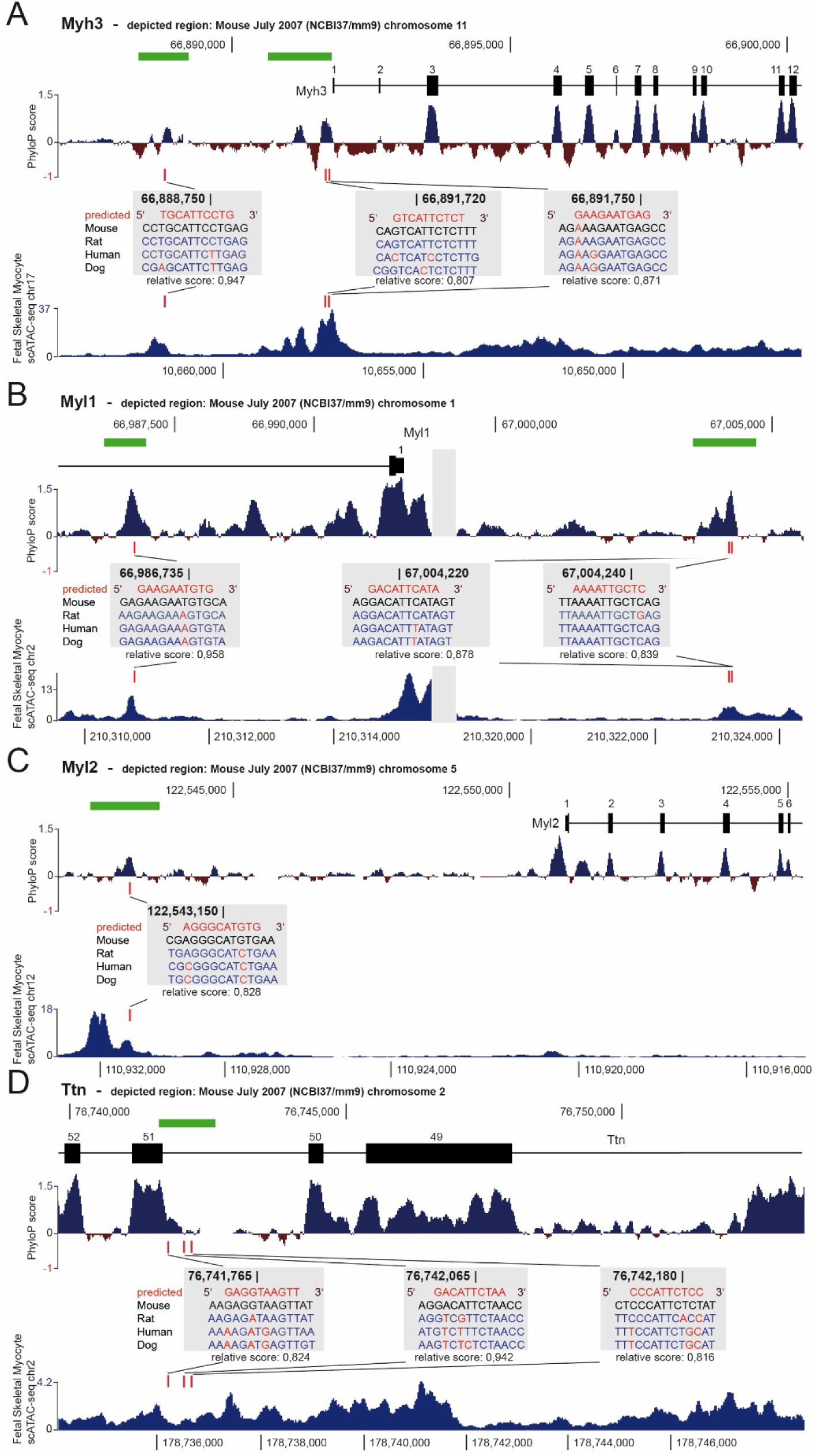
TEAD-bound M-CAT motifs in conserved regions of sarcomere genes Myh3, Myl1, Myl2, and Ttn. Evolutionarily conserved M-CAT motifs for TEAD4 in regulatory regions of the sarcomere genes Myh3 (A), Myl1 (B), Myl2 (C), and Ttn (D). Each panel shows the genomic locus with PhyloP conservation scores, TEAD4 ChIP-seq occupancy peaks in differentiated C2C12 myotubes (bright green), and predicted TEAD binding sites (JASPAR 2018, MA0809.1; score ≥ 80%) highlighted in red boxes together with multispecies sequence alignments. The indicated M-CAT motifs reside within open chromatin regions of human skeletal myocytes as defined by single-cell ATAC-seq. Prediction and overlap analyses were performed as described in the Figure 8 legend.

To functionally test these elements, we cloned genomic fragments containing M-CAT motifs from candidate target genes upstream of a luciferase reporter and compared reporter activity in C2C12 myoblasts versus myotubes. Reporter constructs containing M-CAT-bearing genome DNA fragments from sarcomere genes showed robust activation, particularly in myotubes (Fig. 10A). Enhancer fragments from myotube-formation and differentiation genes also exhibited myotube-biased induction: elements from *Myomaker*, *Myomixer*, and *Myod1* were strongly activated in myotubes, whereas the *Myog* fragment showed less differentiation-dependent responsiveness (Fig. 10B). These results indicate that TEAD-regulated enhancers from both fusion and structural genes display stage-specific activation during myogenic differentiation.

**Figure 10.**
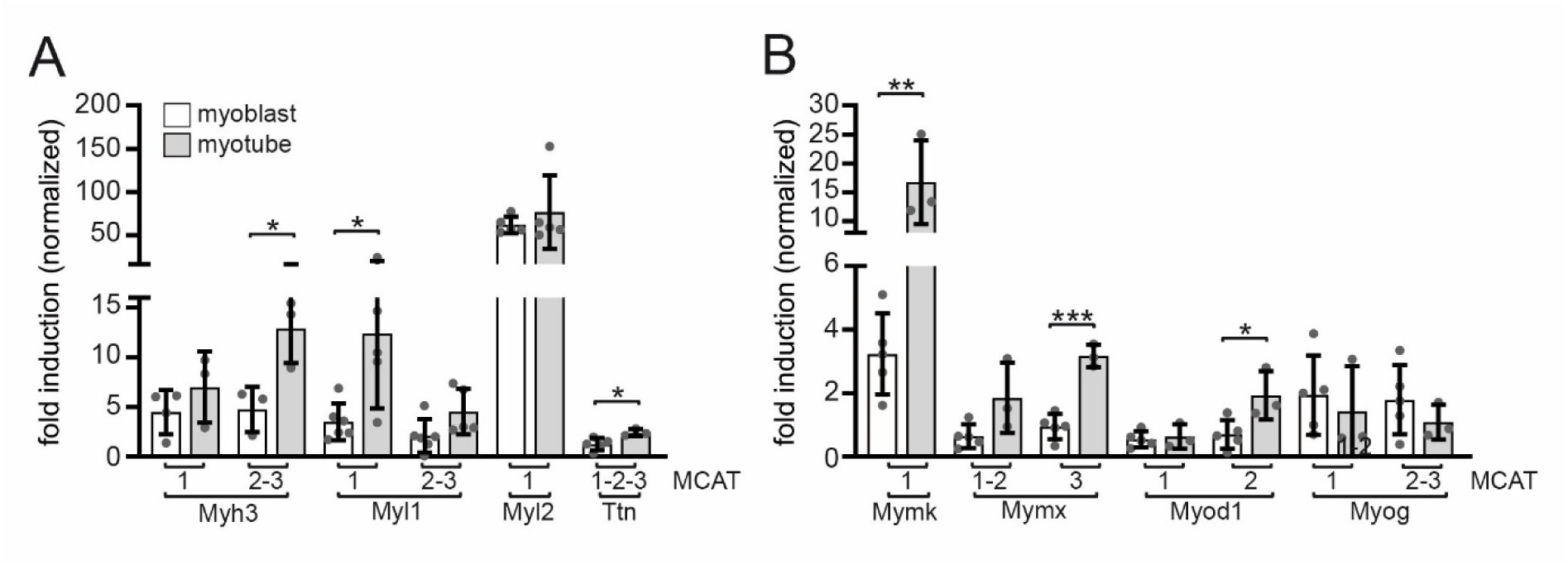
TEAD-binding M-CAT motifs enhance transcription of myogenic genes involved in myotube formation and sarcomere assembly. Genomic regions containing M-CAT motifs (black rectangles in Figures 8 and 9) were cloned upstream of a luciferase ORF in reporter plasmids to assess TEAD-dependent transcriptional activity. Constructs were transfected into proliferating C2C12 myoblasts, and luciferase activity was measured 48 h later (myogeneic genes). Alternatively, transfected myoblasts were differentiated into C2C12 myotubes (day 5) before luciferase assays were performed (sarcomere genes). Graphs summarize obtained luciferase data for sarcomere genes (A), or myogenic genes (B). Data are presented from at least three independent experiments, each performed in triplicate. Bar colors indicate whether assays were performed in myoblasts or myotubes. Statistical significance was assessed using an unpaired Student’s t-test.

Taken together, these findings support a model in which TEAD transcription factors, particularly TEAD4, engage evolutionarily conserved M-CAT motifs near myogenic and sarcomeric genes to promote their expression in differentiating myotubes. By combining TEAD ChIP-seq, motif conservation, chromatin accessibility, and reporter assays, we define a set of TEAD-linked regulatory elements that underlie the myotube-formation and sarcomere gene programs controlled by YAP1/TAZ-TEAD signaling in skeletal muscle.

## DISCUSSION

### Divergent and overlapping roles of YAP1 and TAZ in myogenic maturation

Nuclear YAP1 and TAZ, which lack intrinsic DNA-binding domains, associate with members of the TEAD transcription factor family to activate canonical target genes such as *Cyr61*, *Ctgf*, and *Ankrd1*. While YAP1 and TAZ share several structural domains (26), an important open question has been how their functions diverge within the myogenic lineage and how their shared pathway impacts muscle fiber maturation. Previous work suggested overlapping roles in promoting myoblast proliferation, with TAZ subsequently switching to enhance myogenic differentiation (22), and TAZ-TEAD4 signaling has been implicated in regulating synaptic gene expression at the NMJ (23). Consistent with this, skeletal muscle-specific cDKO mice were reported to die shortly after birth, likely due to severely impaired NMJ formation (23). Our current study demonstrates that this perinatal lethality is accompanied by profound structural defects within the muscle fibers themselves, indicating that intrinsic myofiber abnormalities substantially contribute to the lethal phenotype.

In line with findings in Drosophila flight muscle, where the Yorkie-Scalloped axis tunes sarcomere gene transcription (49), our transcriptomic profiling and high-resolution imaging reveal that YAP1 and TAZ are critical for mammalian sarcomere morphogenesis. While our earlier study (23) using conventional imaging did not detect overt structural alterations, the refined analysis presented here uncovers a markedly disorganized cytoskeleton with misassembled sarcomeres and disturbed Z-disk/M-band alignment in the cDKO mice. These findings position YAP1/TAZ-TEAD signaling as a conserved regulator of sarcomere organization across phyla.

Importantly, reduced hindlimb muscle size appears not merely a secondary consequence of sarcomere defects. Myotube formation and sarcomere assembly are sequential but tightly coordinated processes: myotubes provide the structural and mechanical framework necessary for sarcomeres to mature. Our developmental analysis at E14.5, E16.5, and E18.5 provides temporal resolution of these events (Fig. 3). The data do not support a model in which only the first or second wave of myotube formation is selectively affected; rather, both primary scaffolds and secondary myotubes show impaired growth and maturation. Instead, the loss of YAP1 and TAZ appears to compromise the entire myogenic program: myotubes with reduced CSA are formed initially, and those that do emerge fail to undergo the characteristic radial growth and maturation (Fig. 3G).

This is particularly evident in cDKO SOL muscles at E18.5, where persistent doughnut-shaped fibers lacking central MYH3 signal indicate that even primary scaffolds fail to complete sarcomere assembly (Fig. 3A). This observation is compatible with the idea that a critical mechanical tension state within the myotube is required to initiate periodic myofibrillogenesis. Consistently, *ex vivo* fusion assays using tamoxifen-induced Pax7-CreER^T2^; *Yap1*/*Wwtr1* double-knockout myoblasts revealed significantly reduced myotube formation, indicating that myotube formation is fundamentally less efficient in the absence of both factors (Fig. 5A-C). Future SHG (second harmonic generation) analysis of cDKO cultures could help determine whether myofibrillar architecture is already compromised at the nascent myotube stage, as suggested by previous studies in 3D-engineered muscles (54). Finally, the differential susceptibility of individual muscle groups - where some distal regions are completely absent in the cDKO mice - suggests that factors such as mechanical load, innervation, or fiber-type-specific requirements for YAP1/TAZ signaling may modulate the severity of the phenotype.

### A YAP1/TAZ-TEAD transcriptional program coordinates fusion and sarcomere maturation

To gain mechanistic insight into how YAP1 and TAZ regulate muscle development, we used RNA-seq to identify target genes downregulated in the absence of these co-activators. In line with our histological observations of developmental arrest of myofibers in the cDKO at E16.5-E18.5 (Fig. 3), we identified a coordinated reduction in transcripts essential for both myoblast fusion and sarcomere assembly, and we confirm downregulation of previously (23) proposed TEAD-regulated NMJ genes (Fig. 4F), underscoring that synaptic and structural programs are jointly controlled. Notably, the fusion-critical gene Mymk showed significantly reduced levels (Fig. 4F), providing a molecular explanation for the sparse myotube formation observed in our Pax7-CreER^T2^-derived cultures (Fig. 5A). Furthermore, the downregulation of key structural genes - including *Myh3*, *Myl1*, *Myl2*, *Ttn*, and members of the palladin/myomesin families - corroborates the failure of sarcomere assembly and the persistent doughnut-like morphology seen in cDKO primary myotubes.

By correlating previously reported TEAD1/4 ChIP-seq data (34) with our cDKO neonate muscle transcriptomes, we found that a large proportion of downregulated genes lie near TEAD binding peaks and are associated with muscle contraction. This strongly suggests that YAP1/TAZ-TEAD protein complexes directly control the muscle structural core at the transcriptional level. Using phylogenetic conservation and scATAC-seq data, we identified conserved M-CAT motifs in open chromatin regions of human myocytes that coincide with reported TEAD4 peaks. The functional relevance of these M-CAT motifs, particularly for candidates such as *Myh3*, *Myl1*, *Myl2*, and *Ttn*, was further confirmed by reporter assays, mirroring our previous findings on postsynaptic gene regulation (23).

Our findings integrate well into the emerging framework of muscle development, in which a MYOG-KLF5-TEAD4 transcriptional complex has been proposed to activate myogenic gene programs (35). We extend this model by showing that YAP1/TAZ are required co-activators of TEAD-dependent myogenic programs that drive efficient myotube formation and sarcomere maturation *in vivo*. Specifically, the identification of an M-CAT motif within the recently described 4.23-kb Myh3 regulatory element (55,56) highlights a direct molecular link between Hippo signaling and thick-filament gene regulation. Beyond these shared transcriptional programs, analysis of viable single knockout mice definitively confirms that YAP1 and TAZ are not interchangeable in skeletal muscle (Suppl. Fig. 2). In adult *Wwtr1* knockout SOL muscles, we observed increased fiber CSA and a slow type I fiber bias (Suppl. Fig. 2B, D), suggesting that TAZ may normally act to restrain or fine-tune the timing of differentiation and fiber-type specification. In contrast, *Yap1* deficiency resulted in reduced CSA across all fiber types (Suppl. Fig. 2E), consistent with earlier reports assigning YAP1 a role in supporting early myogenic expansion (23,36,56). The distinct adult phenotypes—hypertrophy in Wwtr1 mutants versus hypotrophy in Yap1 mutants—underscore that YAP1 and TAZ are not interchangeable even within the same pathway, but instead tune fiber growth and type composition through separable functions.

Furthermore, our data point to a sophisticated signaling crosstalk. The reduction of Axin2 and Ctnnb1 levels in the mutants (data not shown) is consistent with a functional interaction between YAP1/TAZ and canonical Wnt signaling (33,57). Potential interactions with TGF-β signaling may likewise contribute to the observed defects in *Mymk* expression and fusion. These pathways represent key avenues for future investigation to dissect how Hippo signaling integrates extrinsic cues with the myogenic transcriptional machinery.

In summary, our data indicate that the Hippo pathway, acting through YAP1/TAZ-TEAD protein complexes, provides critical temporal and quantitative control of sarcomere gene expression. By coordinating the transcriptional programs for myotube fusion with those for myofibril assembly, this pathway synchronizes muscle fiber growth with the precise morphogenesis of its contractile apparatus - a process that is fundamentally disrupted upon the combined loss of YAP1 and TAZ.

## DATA AVAILABILITY

The data underlying this article are uploaded in the Gene Expression Omnibus (GSE270019) at https://www.ncbi.nlm.nih.gov/geo/. All further data generated or analyzed during this study are included in this published article and its supplementary files.

## FUNDING

BvE was supported by the Wilhelm Sander-Stiftung [2022.084.1], the Federal Ministry of Education and Research (BMBF) [16GW0271K], the German Research Foundation (DFG) [EY 120/4-1], and Deutsche Krebshilfe (German Cancer Aid). T.P. was funded by the Polish National Science Centre [2020/37/B/NZ3/03909]. S.H. was supported by the Interdisciplinary Centre for Clinical Research at the University Hospital Erlangen (IZKF ELAN P185) and by the German Research Foundation (DFG) [HA 3309/3-4].

## ACKNOWLEDGMENTS

We gratefully acknowledge the support of the Core Facility Next Generation Sequencing at the FLI for library preparation and sequencing, in particular Ivonne Goerlich and Marco Groth.

## Author contributions

LG and SH performed cellular and animal experiments, cell culture studies, statistical analyses, histological analyses, and analysis of experimental data. SH conceived and directed the research and supervised interpretation of the experimental data. TP organized and analyzed the electron microscopy data. MS analyzed and evaluated pathological impairments in skeletal muscles. BvE performed transcriptome analysis and all other bioinformatic analyses. LG analyzed evolutionarily conserved regions (ECRs) and ATAC-seq correlations for candidate targets. LG, BvE, and SH wrote the manuscript.

## CONFLICT OF INTEREST DISCLOSURE

The authors declare that they have no conflicts of interest.

## SUPPLEMENTARY DATA

**Supplementary FIGURE 1:**
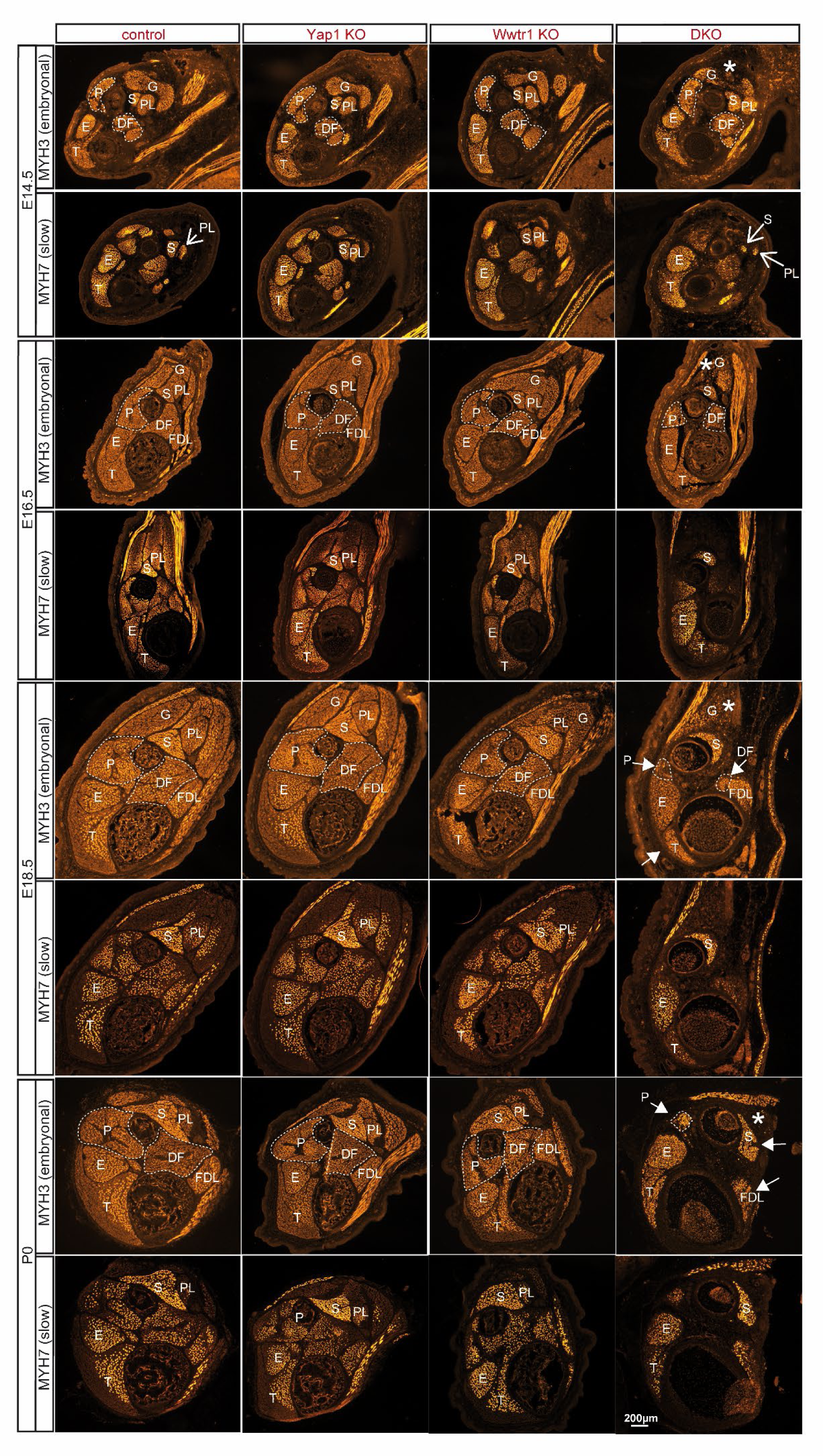
Developmental time-course of hindlimb muscle architecture in control, single knockout, and cDKO embryos and newborns. Representative cross-sections of embryonic hindlimbs from control, Yap1 knockout, Wwtr1 knockout, and *Yap1*/*Wwtr1* conditional double knockout (cDKO) mice at E14.5, E16.5, E18.5, and P0. Sections were stained for MYH3 (embryonic) or MYH7 (slow) as indicated. Identifiable muscle groups are labeled: G, gastrocnemius; S, soleus; P, peroneus muscle group; FDL, flexor digitorum longus; E, extensor digitorum longus; PL, plantaris; DF, deep flexor muscle group. White asterisks mark the gastrocnemius muscle, which is markedly reduced in cDKO embryos. Solid white arrows indicate prominent structural abnormalities in cDKO samples. In the DKO, several muscles disappear completely, e.g., plantaris, distal parts of tibialis (compare MYH3 stained tibialis control with cDKO), change their morphology and position (e.g., soleus); and show disrupted muscle architecture (e.g., gastrocnemius, where the medialis does not form and disappears completely by E18.5). Due to space, open white arrows label individual muscle groups across genotypes. At E18.5, solid white arrows do both, label muscle and point to structural abnormalities. For orientation, DF and P muscle groups are outlined with white dashed lines in the MYH3-stained sections. Apparent reductions in SOL and plantaris size at E14.5 in cDKO MYH7-stained images are most likely due to sectioning angle. Scale bar, 200 μm.

**Supplementary Figure 2.**
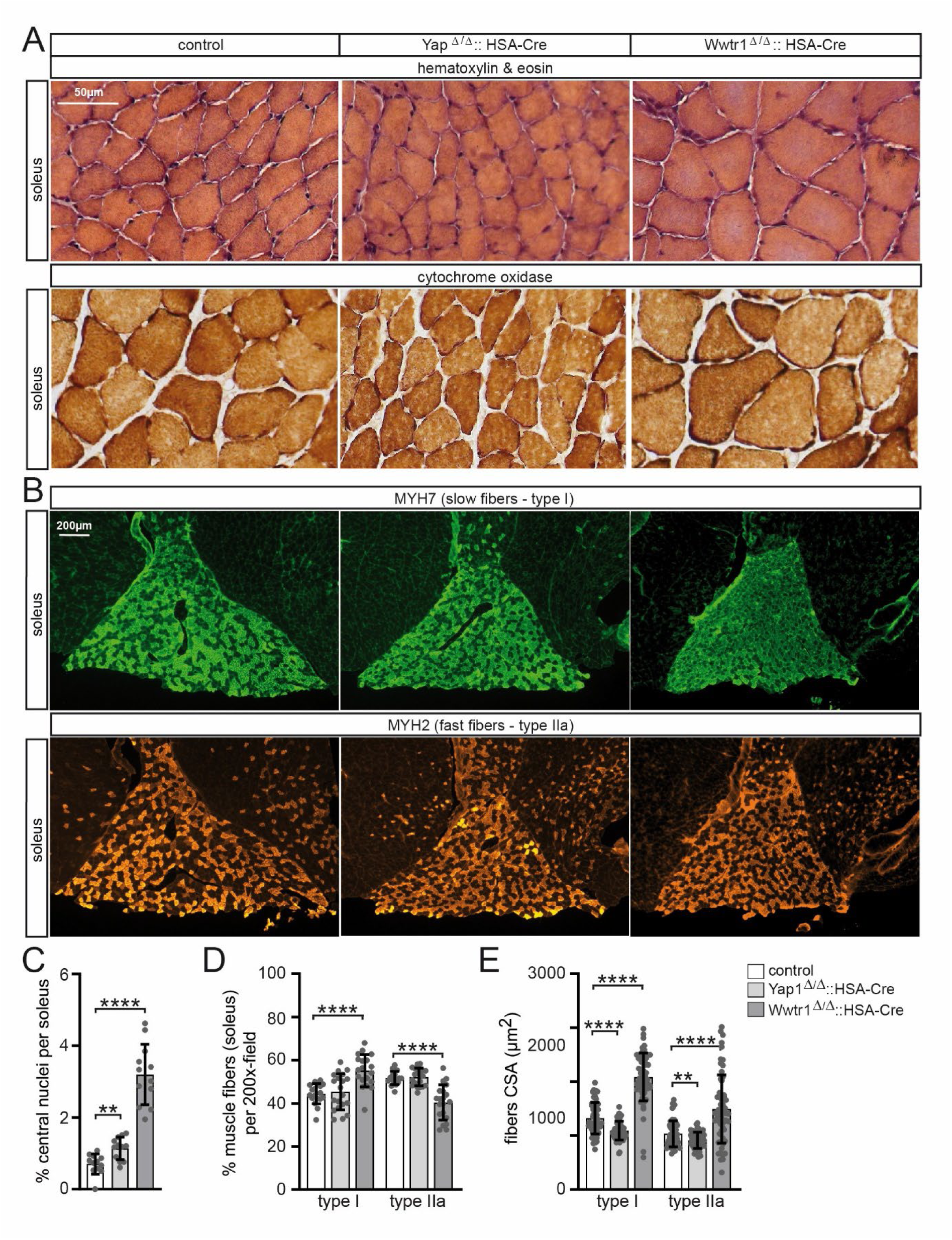
Histological and fiber-type analysis of skeletal muscle in control, Yap1, and Wwtr1 single knockout adult mice. (A) Representative transverse sections of adult soleus muscle stained with hematoxylin and eosin or for cytochrome oxidase activity. Both single knockout models display normal overall muscle architecture compared with controls. Scale bar, 50 µm. (B) Immunofluorescence staining of soleus muscle for MYH7 (type I, slow fibers; green) and MYH2 (type IIa, fast fibers; orange/red). Fiber-type distribution and organization appear comparable between genotypes. Scale bar, 200 µm. (C) Quantification of centrally nucleated fibers per soleus muscle. (D) Percentage of type I and type IIa fibers per 200× field in soleus muscle. (E) Cross-sectional area (CSA) of type I and type IIa fibers. Data in C-E are presented as mean ± SD; individual data points represent biological replicates. N=4 mice per genotype were analyzed. Statistical significance was assessed by unpaired Student’s t-test.

**Supplementary table 1.** Tabular presentation of used BTX, primary and secondary antibodies for immunostaining.

| Target / Name | Species | Antibody reference | Supplier | Dilution |
| --- | --- | --- | --- | --- |
| MyH7 (MyH-slow) | mouse IgG2B | BA-F8 | DSHB | 1:1000 |
| MyH2 (MyH-IIa) | mouse IgG1 | SC-71 | DSHB | 1:1000 |
| MyH3 (MyH-emb) | mouse IgG1 | F1.652 | DSHB | 1:200 |
| MyH slow | mouse IgG1 | M8421 | Sigma | 1:200 |
| MyH fast | mouse IgG1 | M4276 | Sigma | 1:200 |
| rhodamine- conjugated α-bungarotoxin (BTX) | - | T1175 | Invitrogen | 1:2500 |
| anti-mouse IgG2b Alexa488 | goat | A-21141 | Invitrogen | 1:1000 |
| anti-mouse IgG1 Alexa546 | goat | A-21123 | Invitrogen | 1:1000 |

**Supplementary table 2.** Summary of identified TEAD1- and TEAD4-occupied sites in the vicinity of sarcomer genes Myl1, Myl2, Myh3, Ttn and myogenic genes Mymk, Mymx, MyoD and MyoG in C2C12 cells at day 0 and day6 of differentiation from (34).

| TEAD1 - day 0 |  |  |  |
| --- | --- | --- | --- |
| *gene* | *chromosome* | *start* | *end* |
| - | - | - | - |
| TEAD1 - day 6 |  |  |  |
| *gene* | *chromosome* | *start* | *end* |
| - | - | - | - |
| TEAD4 - day 0 |  |  |  |
| *gene* | *chromosome* | *start* | *end* |
| - | - | - | - |
| TEAD4 - day 6 |  |  |  |
| *gene* | *chromosome* | *start* | *end* |
| Myl1 | chr1 | 66986211 | 66986969 |
|  | chr1 | 67003571 | 67004722 |
|  | chr1 | 67010900 | 67011898 |
| Myl2 | chr5 | 122542445 | 122543691 |
| Myh3 | chr11 | 66888289 | 66889206 |
|  | chr11 | 66890628 | 66891789 |
|  | chr11 | 66917944 | 66918600 |
|  | chr11 | 66920195 | 66920980 |
| Ttn | chr2 | 76741610 | 76742632 |
| Mymk | chr2 | 26930924 | 26931842 |
| Mymx | chr17 | 45738717 | 45739701 |
|  | chr17 | 45752676 | 45753939 |
| MyoD | chr7 | 53607821 | 53608972 |
|  | chr7 | 53626260 | 53627124 |
| MyoG | chr1 | 136173278 | 136174448 |
|  | chr1 | 136180715 | 136182004 |

**Supplementary table 3.** Full list of identified putative TEAD4 binding sites in TEAD4-occupied regions in the vicinity of sarcomeric genes *Myl1*, *Myl2, MyH3, Ttn* and myogenic genes *Mymk*, *Mymx*, *MyoD* and *MyoG* in C2C12 cells according to ChIP-Seq data in (34). Genomic coordinates refer to Mouse July 2007 (NCBI37/mm9) genome assembly. Note, all sites are located in regions occupied by TEAD4 exclusively in differentiated C2C12 cells.

| Full list of putative TEAD4 binding sites |  |  |  |  |  |  |  |
| --- | --- | --- | --- | --- | --- | --- | --- |
| *gene* | # | *strand* | *sequence* | *chromosome* | *start* | *stop* | *score* |
| Myl1 | 1 | - | CACATTCTTC | chr1 | 66986730 | 66986739 | 0.958 |
|  | 2 | + | GACATTCATA | chr1 | 67004218 | 67004227 | 0.878 |
|  | 3 | + | AAAATTGCTC | chr1 | 67004234 | 67004243 | 0.839 |
| Myl2 | 1 | - | CACATGCCCT | chr5 | 122543147 | 122543156 | 0.828 |
| Myh3 | 1 | + | TGCATTCCTG | chr11 | 66888745 | 66888754 | 0.947 |
|  | 2 | + | GTCATTCTCT | chr11 | 66891719 | 66891728 | 0.807 |
|  | 3 | - | CTCATTCTTC | chr11 | 66891743 | 66891752 | 0.871 |
| Ttn | 1 | - | AACTTACCTC | chr2 | 76741763 | 76741772 | 0.824 |
|  | 2 | + | GACATTCTAA | chr2 | 76742061 | 76742070 | 0.942 |
|  | 3 | + | CCCATTCTCT | chr2 | 76742175 | 76742184 | 0.816 |
| Mymk | 1 | - | AGCATTCCAT | chr2 | 26931259 | 26931268 | 0.968 |
| Mymx | 1 | + | AGCTTTCTTC | chr17 | 45738774 | 45738783 | 0.822 |
|  | 2 | - | AATATACCAG | chr17 | 45739154 | 45739163 | 0.813 |
|  | 3 | - | CACATTCTGT | chr17 | 45753134 | 45753143 | 0.940 |
| MyoD | 1 | - | GGCCTTCCTC | chr7 | 53608896 | 53608905 | 0.837 |
|  | 2 | + | AACATTCCTG | chr7 | 53626745 | 53626754 | 0.965 |
| MyoG | 1 | - | TGCATTCCTG | chr1 | 136173629 | 136173638 | 0.947 |
|  | 2 | + | CAGATTCCAG | chr1 | 136181337 | 136181346 | 0.889 |
|  | 3 | + | AAGATTCTTG | chr1 | 136181303 | 136181312 | 0.834 |

**Supplementary table 4.**
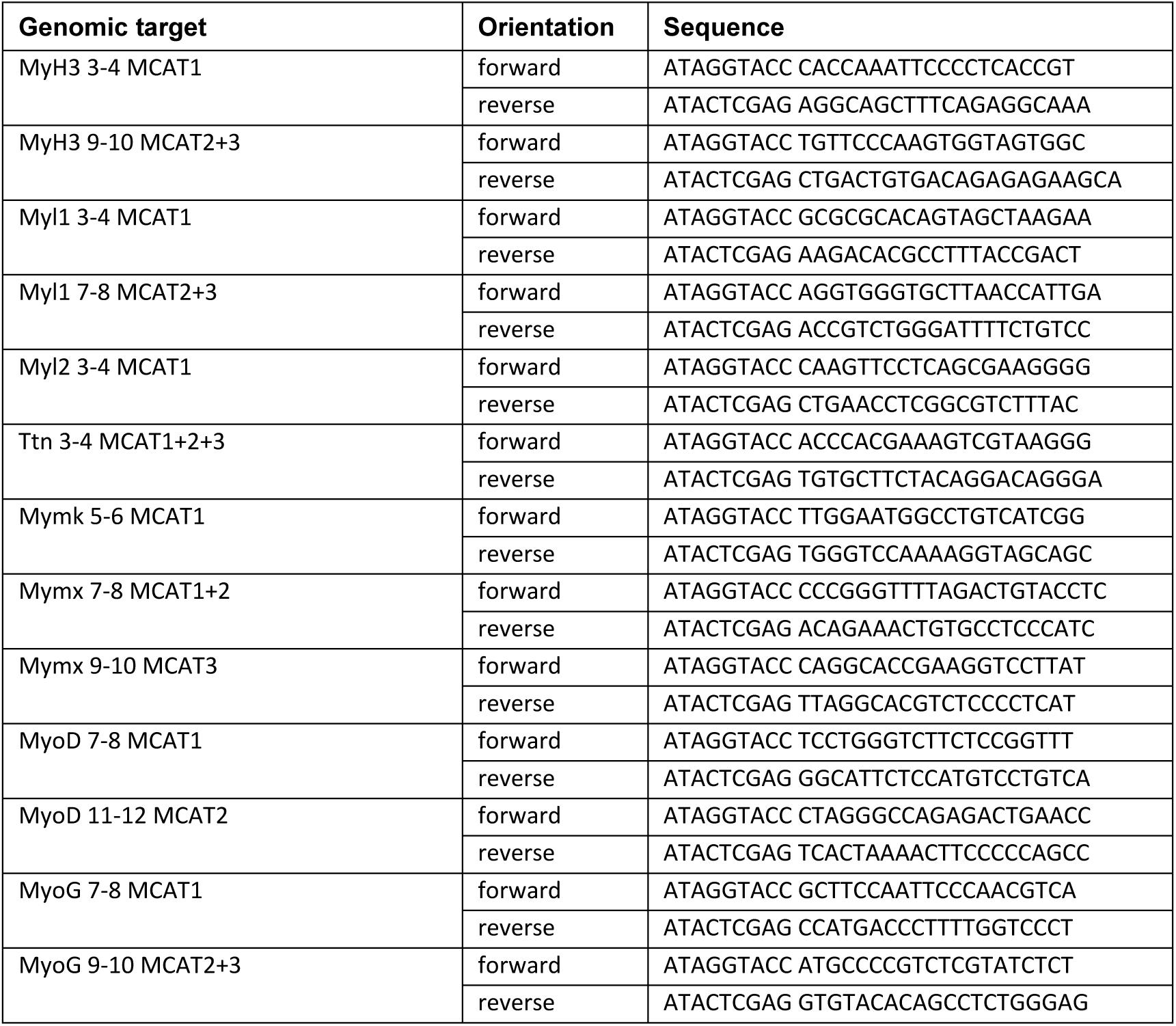
Tabular presentation of oligonucleotide sequences.

## Notes

### Competing Interest Statement

The authors have declared no competing interest.

### Summary of Updates

This version substantially expands the original manuscript with additional experiments and analyses addressing skeletal muscle development, myotube formation, and sarcomere organization. New developmental analyses examine hindlimb muscle architecture and myofiber growth and maturation at embryonic stages E14.5, E16.5, and E18.5 and at birth. Measurements of neonatal hindlimb bones have also been added to distinguish impaired muscle development from altered bone growth. New ex vivo experiments use satellite-cell-derived primary myogenic cultures with inducible Pax7-CreERT2-mediated deletion of Yap1 and Wwtr1. These experiments assess myotube formation and myonuclear incorporation following combined loss of YAP1 and TAZ. The regulatory analyses have been expanded to include candidate elements associated with myogenic and fusion genes, including Mymk, Mymx, Myod1, and Myog, alongside sarcomere genes. Published TEAD1 and TEAD4 chromatin occupancy data are integrated with transcriptomic data, developmental single-nucleus gene expression data, sequence conservation, and chromatin accessibility information. New luciferase reporter assays assess the activity of selected candidate regulatory elements in myogenic cells. The title, abstract, Introduction, Results, and Discussion have been revised to emphasize the relationship between myoblast fusion, myofiber maturation, and sarcomere assembly. Figures have been reorganized and expanded, and the Methods, figure legends, references, and supplementary material have been updated accordingly.

